# Insights into goatpox virus and sheeppox virus genomes from pangenome graphs

**DOI:** 10.64898/2026.03.28.714820

**Authors:** Tim Downing

## Abstract

The capripoxviruses comprise three species: goatpox virus (GTPV), sheeppox virus (SPPV) and lumpy skin disease virus (LSDV). They are large double-stranded DNA viruses with highly conserved core genomes and variable terminal regions. Previous studies have described variation in *Capripoxvirus* gene content, their broader population structure and the contribution of non-coding and structural variation remains opaque. This study investigated the genomic diversity and evolutionary history of GTPV and SPPV using phylogenetics, pangenome variation graphs (PVGs), and gene-specific analyses. We found clear differences in population structure between the two viruses. GTPV had three deeply divergent and genetically stable lineages with limited evidence of recent gene flow, whereas SPPV had weaker clade separation consistent with an ancestral bottleneck followed by recent population expansion. PVG-based analyses indicated that GTPV has a comparatively closed pangenome, while SPPV’s was open, particularly at the inverted terminal repeats (ITRs). Structural and haplotype variation was concentrated at these ITRs, which moderate host immunity and specificity. In several lineages, extended putative ORFs spanning adjacent ITR genes were observed, indicating recurrent structural plasticity at these regions. Patterns of gene-specific conservation and divergence highlighted loci under strong constraint and lineage-specific structural changes that may contribute to host specificity. Together, these results demonstrated how graph-based genome models complement gene-based analyses in resolving poxvirus genome evolution and provide a resource for improved comparative and population genomic studies of large DNA viruses.

**Significance:** The capripoxviruses are economically important livestock pathogens, yet the genomic mechanisms underlying their diversification and host specificity remain poorly resolved. By applying pangenome variation graphs alongside phylogenetic and gene-level analyses, this study reveals fundamental differences in how goatpox and sheeppox viruses have evolved. Goatpox virus had a deeper, more stable lineage structure, whereas sheeppox virus was more recent and diverse. Importantly, structural variation at the inverted terminal repeats emerged as a major driver of genomic diversity, including lineage-specific haplotypes and variable gene structures. These findings demonstrated the value of graph-based genome representations for resolving complex variation in large DNA viruses and provides approaches for improving genomic surveillance, comparative analyses, and future investigations into host range, virulence and tropism.

## Introduction

Goatpox virus (GTPV) and sheeppox virus (SPPV) belong to the *Capripoxvirus* (CaPV) genus in the family *Poxviridae* and primarily infect goats and sheep as hosts, respectively. The other species in the CaPV genus is the lumpy skin disease virus (LSDV) whose preferential host is cattle. Their International Committee on Taxonomy of Viruses (ICTV) names are Capripoxvirus goatpox, Capripoxvirus sheeppox and Capripoxvirus lumpyskinpox, respectively. Collectively, the CaPV are re-emerging transboundary dsDNA viruses that impair animal health, and consequently cause significant economic harm (Casal et al 2018, Vinitchaikul et al 2023). They are all notifiable diseases according to the World Organization for Animal Health (WOAH) (WOAH 2010) because of their wide geographic spread and their economic importance. GTPV and SPPV are endemic across parts of Africa, the Middle East, and Asia, and phylogenetic analyses identify geographically associated sub-clades in these regions. Additionally, SPPV has deep ancestry within Europe dating back several centuries (Binois-Roman et al 2025), with possible replacement by Clade 3.3 (also called clade A1), which has spread into southern Europe in recent years (§an et al 2024). Consequently, the potential for GTPV and SPPV to cause disease outbreaks is a global challenge in relation to animal welfare, economic stability and transboundary spread.

GTPV and SPPV can survive in the environment for long periods of time, and can transmit via contact, unlike LSDV that typically needs an insect vector (Sprygin et al 2025). GTPV can infect cattle and sheep as well, though less effectively (Bhanuprakash et al 2010, Ramakrishnan et al 2017). In the same way, SPPV can experimentally infect goats and cattle, and LSDV goats and sheep. There is vaccine cross-protection between GTPV and SPPV contrasting with the partial protective effect of LSDV vaccination against SPPV, and likewise for SPPV vaccination against LSDV (Hamdi et al 2010). This has resulted in the wider use of GTPV as a CaPV vaccine component thanks to its broader cross-protective profile, such as reducing the risk of LSDV infection (Haegeman et al 2025).

CaPV possess a linear genome with lengths of about 150 Kb, are AT-rich (75%), and typically have ∼147-156 orthologous genes (Tulman et al 2001). The estimated interspecies genome-wide similarity is about 96%, stemming from the ancient diversification of LSDV, GTPV and SPPV (Tulman et al 2002). All share diverse inverted terminal repeats (ITRs) at both genome termini whose genes moderate host range, interactions and virulence (Kara et al 2003, Zhang et al 2025). These ITRs form complementary hairpin structures at both ends, which are essential for concatemer resolution and the formation of mature genomes during replication (Shenouda et al 2022). These ITRs are also structurally dynamic, frequently exhibiting deletions, duplications, and rearrangements that are associated with attenuation, host adaptation, and vaccine derivation. The 5’ (LSDV ORFs 1-23) and 3’ (LSDV ORFs 124-156) are more variable and typically support functions linked to immunomodulation and virulence. The intervening genomic region (LSDV ORFs 24-123) is highly conserved and encodes core replicative and structural machinery (Tulman et al 2001, Tulman 2002, Haga et al 2024). In addition, there are nine genes in LSDV that are pseudogenised or truncated in GTPV and SPPV that moderate virulence and host range (Tulman et al 2002).

The genome evolution of CaPV remains understudied, particularly for GTPV and SPPV. To address this, this study uses pangenome variation graphs (PVGs) (also known as pangenome graphs or sequence graphs) to illuminate genomic diversity and the evolution of GTPV and SPPV in all available high-quality genome assemblies (Downing 2025). By representing multiple genomes as paths through a shared graph structure, PVGs enable the joint analysis of SNP, indel and structural variation (Eizenga et al 2020, Garrison & Guarrcino 2023). Here, a gene-level analysis of genome-wide diversity in both species was created to probe their population structure and genetic variability, including the ITRs. Additionally, this study linked GTPV and SPPV evolution to that of LSDV using a consistent nomenclature throughout based on the Oman LSDV reference genome (Wright et al 2026) to provide a consistent framework for genes that are intact in LSDV but altered in GTPV and SPPV. Lastly, representative PVGs were constructed for both species, with potential to improve mutation detection during read mapping of short read CaPV sequencing libraries. Together, this work provided a graph-based perspective on GTPV and SPPV genome evolution and highlighted how differences in population structure, terminal genome plasticity, and lineage-specific variation may contribute to host specificity and virulence.

## Material and Methods

### Genome assembly collection, alignment and phylogenetic analysis

All available complete and valid GTPV (n=14) and SPPV genomes (n=30) were downloaded from the NCBI nucleotide database (Sayers et al 2024) (5^th^ August 2025) (Table S1). KT438551.1 and KT438550.1 were also two listed SPPV complete genomes, but were excluded because their genome assembly quality was low.

This initial dataset was aligned with Mafft v7.453 (Katoh & Standley 2013) using automatic optimisation and default parameters. Phylogenetic reconstruction of the samples’ evolutionary relationships was conducted using RAxML-NG (Randomised Axelerated Maximum Likelihood) v1.2.0 (Kozlov et al 2019) with a GTR (general time reversible) model and gamma substitution rate heterogeneity selected as the best-fit model by modeltest-ng (Darriba et al 2020). Bootstrap support was calculated using both Felsenstein Bootstrap Proportion (FBP) and Transfer Bootstrap Expectation (TBE) methods with 1000 replicates where support values ≥90% were displayed. These phylogenies were mid-pointed rooted and visualised using R v4.3.2 (R Core Team 2024) packages ape v5.8-1 (Paradis & Schliep 2019), phangorn v2.12.1 (Schliep 2011), treeio v1.28.0 (Wang et al 2020), ggtree v3.12.0 (Yu et al 2017) and ggplot2 v2_3.5.2 (Wickham 2016). Principal components analysis (PCA) was implemented using R based on the genome-wide SNP data. As above, the corresponding sequences for each gene were extracted, aligned with mafft, their phylogenies were reconstructed with RAxML. PCA was implemented to detect population structure patterns not apparent in the phylogenies.

### Pangenome variation graph creation, analysis and annotation

Panalyze v1.0 (Tennakoon et al 2025) implemented PVG construction, analysis and annotation of these GTPV (n=14) and SPPV (n=30) genomes. Panalyze created PVGs in Graphical Fragment Assembly (GFA) format using pangenome graph builder (PGGB) v0.4.0 (Garrison et al 2024) and wfmash v0.7.0 (Guarracino et al 2021), with Multiqc v1.14 (Ewels et al 2016) to collate and assess PVG metrics. Panalyze analysed these PVGs using ODGI (optimized dynamic genome/graph implementation) v0.8.3 (Guarracino et al 2022) to extract SNPs, indels and compound mutations using VG from the PVG paths (Garrison et al 2018). Subgraphs were selected using gfatools v0.4 (Li 2023), evaluated with gfastats v1.3.6 (Formenti 2023), analysed with gfastats v0.1 (Formenti 2023), and visualised with VG (Garrison et al 2018). Panalyze visualised the PVGs with ODGI (Guarracino et al 2022) and ggplot2 v2_3.4.4 (Wickham 2016). Panalyze used gfautil v0.3.2 (Kubica 2023) to get mutation rates, coordinates for samples at each path and site, and VCF files. Mutation density and population genetic statistics were computed for the 5’ end (100-13,850 bp), core (13,851-106,910 bp) and 3’ end (106,910 to 150,000 bp) as per previous work (Haga et al 2024).

The PVGs were annotated using Prokka (Seeman 2014) and indexed with SAMtools v1.19 (Danecek et al 2021). Using the reference genome annotation for NC_004002 and NC_004003, BUSCO v5.8.0 (Seppey et al 2019) showed all GTPV and SPPV had all 18 expected BUSCO poxvirus genes. The annotated GTPV, SPPV and LSDV genomes were aligned and compared using NC_004003, NC_004002 and PV877838 (Wright et al 2026) to extract gene boundaries for CDSs present in LSDV that were absent in GTPV and SPPV (Table S2). The Prokka GFFs were converted to gene transfer format (GTF) with gffread (Pertea & Pertea 2020) and parsed. The annotation position information was merged with the ODGI (Guarracino et al 2022) rendering of the PVGs, creating one CSV annotation file per PVG, which was visualised using Bandage v0.8.1 (Wick et al 2015).

### Pangenome size and growth estimation

Panalyze inferred PVG growth using a sequence-based approach with Panacus v0.2.3 using default coverage thresholds that included all segments (Parmigiani et al 2024a), and a k-mer-based method in pangrowth (Parmigiani et al 2024b). Both estimated the total number of mutations (*n*) in a sample collection (*N*) for an estimated parameter *gamma* as: *n*□*=*□*ĸN*^*gamma*^, where *gamma* reflected a closed PVG if negative and an open PVG if positive based on Heap’s Law, and *ĸ* was a fitting parameter. They also estimated the first derivative of this, the rate of new mutations per sample (Δ*n*)□as: Δ*n =*□*kN*^*-alpha*^, which is based on *alpha<1* denoting an open PVG and *alpha>1* a closed one, and *k* was a fitting parameter (Downing & Decano 2019). This modelled the rate of addition of new nodes or k-mers as each sample was added to the PVGs. Here, a new nodes or new k-mer indicated a new mutations Pangrowth also estimated the core PVG size based on k-mer differences. Panalyze also used ODGI heaps (Guarracino et al 2022) to infer PVG growth using 1000 simulations.

### Community detection and visualisation

Community detection was performed on PVGs, where these communities reflect the relative similarity of samples based on PVG data. Here, these were identified with Panalyze using wfmash (Guarracino et al 2021) and PGGB (Garrison et al 2023) with a k-mer of 19 bp, a window size of 67 bp, a threshold of 80% sequence mash-level identity, and ten mappings per node. Panalyze evaluated these communities with PGGB scripts (Garrison et al 2023) that built a network with the contigs as nodes and mappings as weighted edges, where the latter were proportional to the mapping’s similarity and length (Guarracino et al 2023). Panalyze visualised these networks with igraph tools (Csardi & Nepusz 2006). The mappings threshold was the number of detected haplotypes minus one for the given region (Guarracino et al 2023).

### Tests for selective processes

We tested for selection using the direction of selective (DoS) metric expressed as: DoS = D_N_/(D_N_+D_S_) - P_N_/(P_N_+P_S_) (Stoletzki & Eyre-Walker 2011) where D_N_ and D_N_ stand for the number of ancestral nonsynonymous and synonymous substitutions (respectively) between the GTPV and SPPV species, and P_N_ and P_S_ stand for the number of current population-level (within-species) nonsynonymous and synonymous mutations (respectively). A DoS value near 0 is consistent with neutrality, DoS < 0 with purifying selection, and DoS > 0 with an excess of nonsynonymous fixed differences consistent with adaptive evolution. This analysis included an existing dataset of 121 LSDV genomes for comparison (Wright et al 2025).

## Results

### Contrasting levels of population structure within goatpox virus (GTPV) and sheeppox virus (SPPV)

Pangenome variation graph (PVGs) were created for available GTPV (n=14) and SPPV (n=30) genomes to explore their genomic diversity (Table S1). GTPV and SPPV had marked differences in their population structures and evolutionary histories, consistent with previous work (L’Hote et al 2026). GTPV had three separate clades named 2.1 (n=6 samples), 2.2 (n=5) and 2.3 (n=3) based on phylogenetic, PCA, network-based interpretation of genome-wide SNPs (Figure 1A). Repeating this for the core region, 5’ end and 3’ end reiterated this overall topology (Figure S1). Here, an existing clade naming system was used (Biswas et al 2019). In GTPV, the first PC (56% of variation) split clades 2.1 and 2.2 from 2.3, and the PC2 (30%) separated all three clades (Figure 1B). This suggested long-term genetic differentiation of these GTPV clades, with no evidence of recent admixture in this dataset (Figure 1C). Clade 2.3 was the only clade without vaccine-related specimens and contained the sole African sample (MN072624), together with two Middle Eastern isolates. As shown below using LSDV-rooted phylogenies, this clade also had the deepest evolutionary root. A genome-wide comparison of GTPV vaccine-related versus field samples found no SNPs differentiating these groups. With Clade 2.2, comparison of the two vaccine-linked samples against the other three identified 23 differentiating SNPs: 21 at the 5’ ITR near the hairpin sequence, another at LSDV037 (30,904 bp in NC_004003) and one at LSDV066 (56,939 bp).

**Fig 1.**
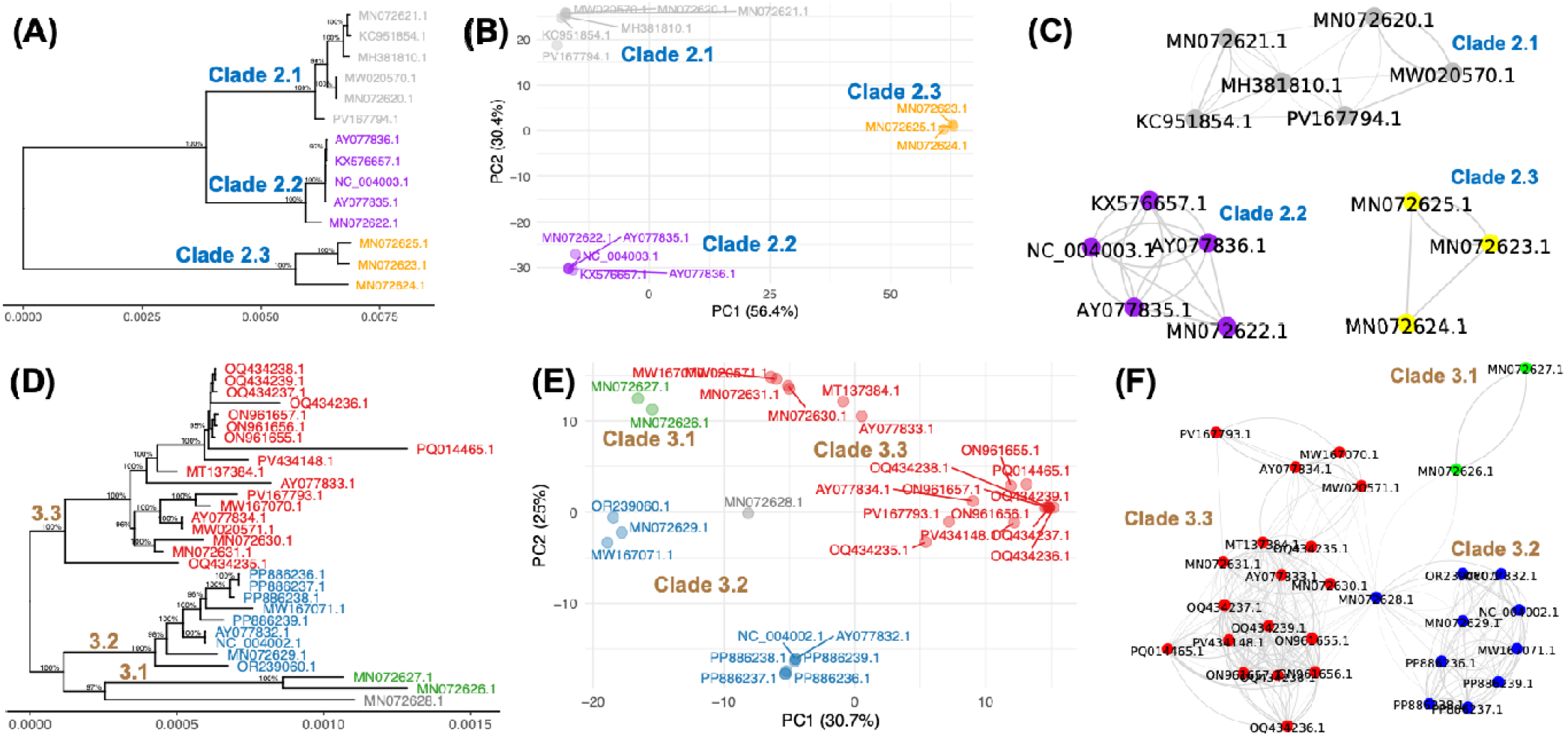
Genome-wide SNP diversity in the GTPV and SPPV PVGs. (A) A midpoint-rooted GTPV phylogeny showing three clades: 2.1 (grey, n=6), 2.2 (purple, n=5) and 2.3 (orange, n=3). The scale bar shows the numbers of substitutions per site. Nodes with bootstrap support >90% are shown. (B) A GTPV PCA where PC1 represented 56% of variation, and PC2 30%. (C) A GTPV network where each node represents one sample. (D) A midpoint-rooted SPPV phylogeny showing three clades: 3.1 (green, n=2), 3.2 (blue, n=9) and 3.3 (red, n=18). (E) A SPPV PCA where PC1 had 31% of variation, and PC2 had 25% of it, and MN072628.1 (grey) was likely a rare lineage. (F) A SPPV network where each node represents one sample.

SPPV had three clades based on the phylogenetic patterns, PCA and networks of PVG data: 3.1 (n=2 samples), 3.2 (n=9) and 3.3 (n=18 samples) (Figure 1D). Within all three SPPV clades, there was a high level of genetic variability that was informed by PCA. Repeating this for the core region, 5’ end and 3’ end reiterated this overall pattern (Figure S2). SPPV PC1 (31%) and PC2 (25%) together showed high diversity within Clade 3.3 (Figure 1E), especially for MW167070, MW020571, MN072630, MN072631, AY077833 and MT137384 that were separate from the main 3.3 cluster. SPPV PC1 and PC2 separated Clade 3.2 into two paraphyletic clusters with three (3.2.1: MW167071, OR239060, MN072629) and six (3.2.2: AY077832, NC_004002, PP886236, PP886237, PP886238, PP886239) samples. Clade 3.1 was genetically distinct and was composed of two vaccine-related samples. Clade 3.3 was spread across Asia and the Middle East. SPPV Clades 3.2 was found in the Middle East, of which subclade 3.2.2 has spread more recently into southern Europe. A genome-wide comparison of SPPV vaccine-related versus field samples found no SNPs differentiating these groups, nor any when this was applied to Clade 3.3 alone.

The sole SPPV genome from Africa (MN072628) was not assigned to any of the other clades. The sample had a central position in the PCA (Figure 1E) and had a long external branch (Figure 1D). MN072628 had similarity to Clade 3.1, four samples from Clade 3.3 (OQ434235, MN072631, MN072630, MW020571.) and four from Clade 3.2 (PP886236, PP886237, PP886239, NC_004002) in the PVG network (Figure 1F). PC3 (10%) and PC4 (6%) separated Clade 3.3 and MN072628, suggesting that it was a rare lineage, which was also supported by simplot analysis showing no notable variation in genome-wide similarity with NC_004002 (Figure S3). This also pointed to a sampling gap and the need for more SPPV genomes from the African continent.

The rooting of these GTPV and SPPV genomes was explored using LSDV sample OQ555660.1, which recapitulated the same GTPV and SPPV phylogenetic structures as previous work (Biswas et al 2019) (Figure S4). This was supported by phylogenetic analysis of a larger sample of (n=116) LSDV genomes combined with GTPV and SPPV ones (Figure S5). This showed that the population structure was weaker and more recent in SPPV, such that the scale of diversity in SPPV was comparable to a GTPV clade. GTPV Clade 2.3 had the most evolutionary deep root, suggesting the differences within GTPV may inform host specificity given that some GTPV isolates infect sheep as well as goats (Biswas et al 2019).

### GTPV and SPPV pangenome variation graph structure, openness and diversity

Although there were fewer GTPV samples, its complete PVG length was almost as long, spanning 160,732 bp compared to SPPV’s 161,135 bp. The GTPV PVG had more nodes than the SPPV PVG (7,801 vs 3,044) and more edges (10,644 vs 4,208). The median PVG length was 147,684 bp for GTPV and 148,183 bp for SPPV, reflecting the median genome size (Figure S6) that was marginally smaller than LSDV. The shared PVG for GTPV was smaller than for SPPV (142,411 bp vs 144,593 bp), implying that if more GTPV genomes were acquired, its shared PVG would shrink, whereas for SPPV it would not.

The distinct GTPV population structure meant it had a closed PVG (alpha=1.76 from Pangrowth), indicating that if more GTPV genomes were sampled, a small number of new mutations would be discovered (Figure S7). These estimates were confirmed with Panacus for GTPV (alpha=1.63) (Figure S8). In contrast, the SPPV PVG was open (alpha=0.69) (Figure S7), suggesting that many new mutations would be found if more SPPV genomes were examined. The Panacus SPPV value was more marginal (alpha=1.01) (Figure S8), suggesting that moderate SPPV sampling may have been achieved. Nonetheless, the lack of samples from African may be limiting for both species.

Representative PVGs for GTPV and SPPV were created using a genome from each main clade (Table 1) using Panalyze (Tennakoon et al 2026). These allow more resolved inference from mapping of read libraries than a linear reference genome (Wright et al 2025). For GTPV, the representative PVG totalled 152,917 bp, with 6,865 nodes and 9,296 edges. It was made from three genomes with 6,247 mutations between them: field sample MN072621 from Clade 2.1 (Biswas et al), vaccine specimen AY077836 from Clade 2.2 (149,673 bp with 155 CDSs) (Tulman et al 2002), and field isolate MN072625 from Clade 2.3 (Biswas et al) (Table 1). NC_004003 (149,599 bp with 153 CDSs) was not used because it lacked sequence corresponding to LSDV155 and LSDV156 at the 3’ end (LSDV001 and LSDV002 were present at its 5’ end).

**Table 1.** The genome used to create representative PVGs for GTPV and SPPV. MN072628.1 was a unique SPPV sample and was not assigned to a clade.

| Species | Accession | Clade | Genome length (bp) | Number of CDSs | Isolation |  |  |
| --- | --- | --- | --- | --- | --- | --- | --- |
|  |  |  |  |  | Year | Country | Type |
| GTPV | MN072621.1 | 2.1 | 150,308 | 156 | 2005 | Vietnam | Field |
|  | AY077836.1 | 2.2 | 149,673 | 155 | 2000 | Kazakhstan | Vaccine |
|  | MN072625.1 | 2.3 | 150,408 | 153 | 1983 | Yemen | Field |
| SPPV | MN072626.1 | 3.1 | 148,534 | 155 | NA | Iraq | Vaccine |
|  | OR239060.1 | 3.2 | 150,441 | 151 | 2017 | United Arab Emirates | Field |
|  | AY077834.1 | 3.3 | 149,612 | 154 | 1994 | Kazakhstan | Vaccine |
|  | MN072628.1 | NA | 149,906 | 149 | 1977 | Nigeria | Field |

The representative SPPV PVG spanned 151,016 bp with 1,611 nodes and 2,235 edges, making it shorter and less diverse than the GTPV PVG. It had four genomes with 1,596 mutations among them (Table 1): vaccine-related sample MN072626 from Clade 3.1 (Biswas et al 2019), field sample OR239060 from Clade 3.2, vaccine-related specimen AY077834. from Clade 3.1 (Tulman et al 2002), and African sample MN072628 to represent less-sampled diversity (Biswas et al 2019). NC_004002 (149,905 bp) was not used because it only had 148 annotated genes, suggesting that some ORFs were missing or mutated.

Genome-wide variability measured based on sequence differences in the PVGs showed higher diversity in GTPV than SPPV, and this was surprisingly higher at CDS regions compared to non-CDS ones (Figure 2). However, both had a shared pattern of higher diversity at the 5’ (<14 Kb) and 3’ (>107 Kb) ends but less variation in the core region, like previous LSDV work (Haga et al 2024) (Table 2). The 5’ and 3’ ends had 23% and 22% more mutations/Kb than the core genome for GTPV, and 83% and 75% more for SPPV (Table 2). Watterson and Wu’s theta per Kb (Watterson 1975) was greater than nucleotide diversity (pi) per Kb genome-wide for both species, and for GTPV Clades 2.1 (38.0 vs 29.0) and 2.2 (28.2 vs 23.5) as well as SPPV Clades 3.2 (12.5 vs 7.6) and 3.3 (19.4 vs 7.4). This pattern was consistent across the genomic regions, resulting in negative Tajima’s D (Tajima 1989) values for GTPV (-2.48) and SPPV (-2.87). This excess of low-frequency variants could be caused by numerous processes, included a recent population expansion.

**Table 2.**
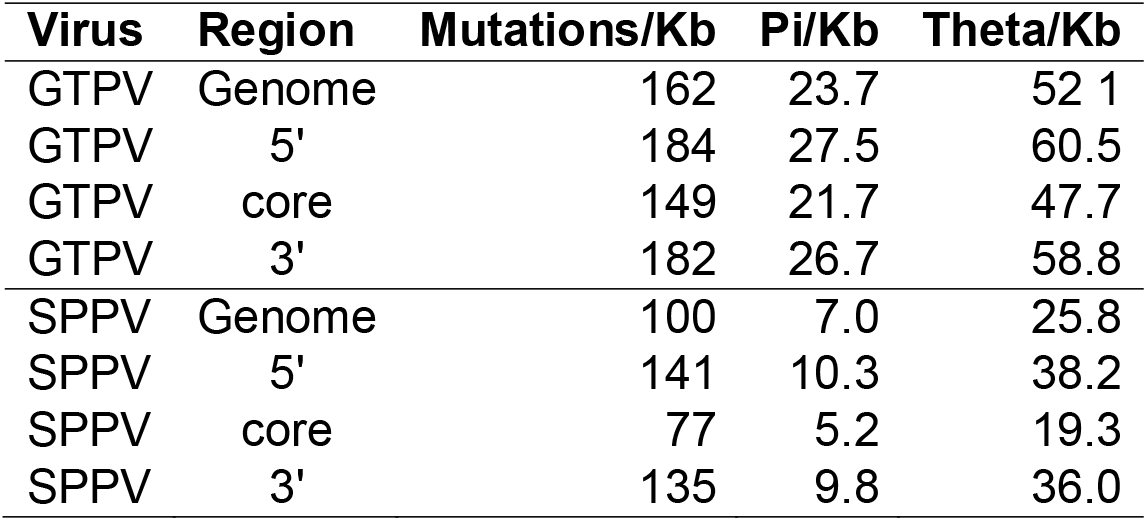
Sequence diversity metrics for GTPV and SPPV. Pi is the nucleotide diversity (the mean number of pairwise SNPs). Theta was calculated as Watterson and Wu’s theta.

**Fig 2.**
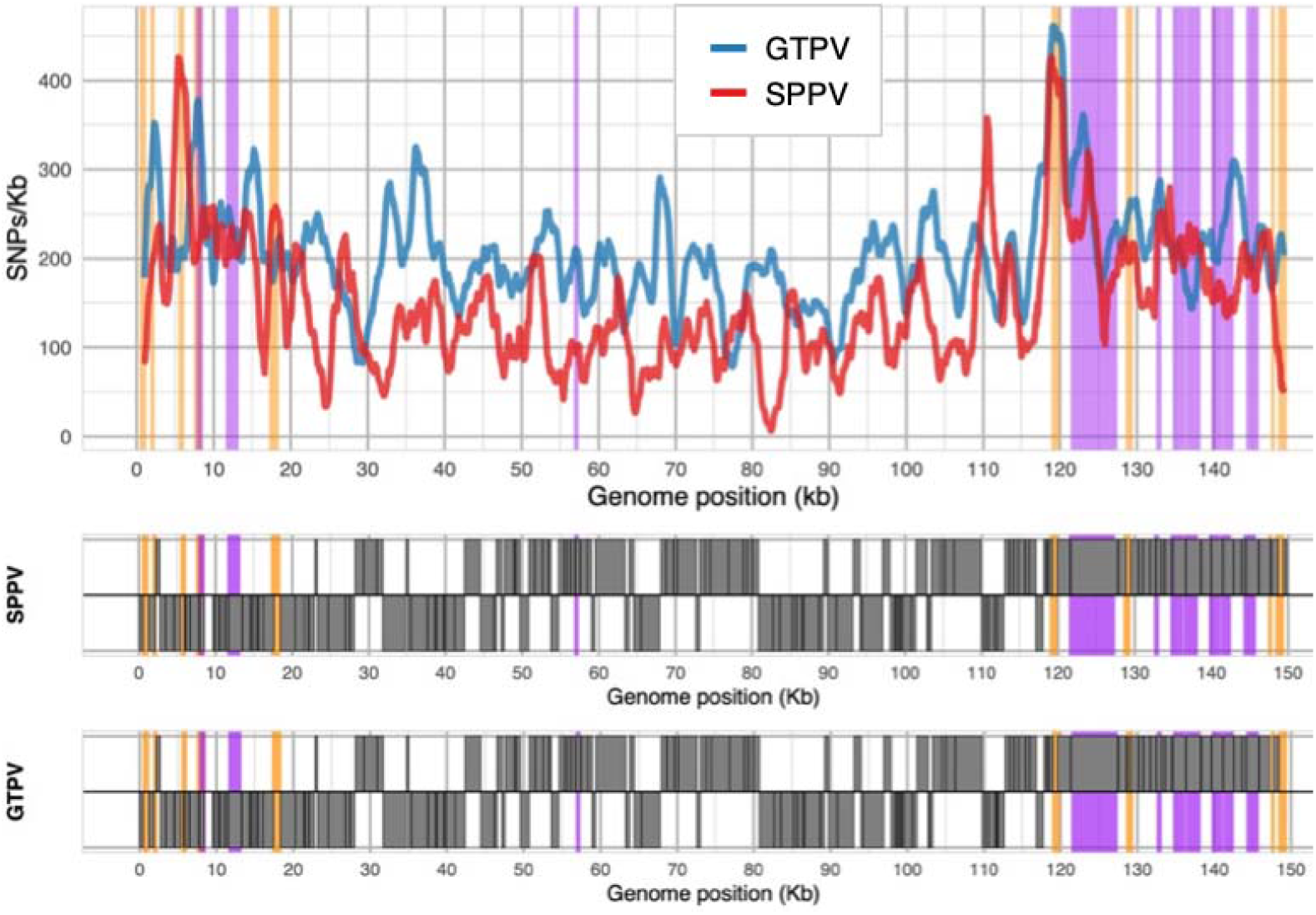
Genome-wide mutation density (mutations/Kb, y-axis) in GTPV (blue) and SPPV (red) highlighting regions associated with CaPV differentiation (LSDV002, LSDV004, LSDV009, LSDV013, LSDV026, LSDV132, LSDV136, LSDV153, LSDV155 in orange) and attenuated virulence (purple) encoding: ankyrin-repeat proteins (LSVD012, LSVD145, LSVD147, LSVD148); kelch-like proteins (LSDV019, LSDV144, LSDV151), a B22R-like protein (LSDV134), or host range factors (LSDV067, LSDV141). The SPPV and GTPV gene annotation across the genomes is represented below the main plot where grey boxes indicated each gene that are read in the forward (lower strip) or reverse (upper strip) directions. The annotation used was from PV877838, NC_004002 and NC_004003.

### Divergent haplotypes at the ITRs highlight novel variation within SPPV

Poxvirus ITRs are repetitive, structurally complex regions underpinning viral replication and virulence (Wright et al 2026) that are also associated with CaPV differentiation (Tulman et al 2002). GTPV and SPPV lack functional LSDV002, LSDV004, LSDV153 and LSDV155 ORFs (Tulman et al 2002). Here, the 5’ ITR of the GTPV PVG had more nodes than SPPV one (226 vs 138) due to its higher diversity (Figure S9). Using LSDV to obtain homologous GTPV and SPPV regions (Table S2), GTPV Clades 2.1 and 2.2 had a 533 bp haplotype at the LSDV002 region (that is similar to MYXV gene M003.2) that differed from a rarer Clade 2.3 487 bp haplotype (Figure 3). Two clade 2.3 samples (MN072624 and MN072625) had truncated LSDV004 ORFs, and MN072625 had a truncated LSDV153 as well. Despite this, LSDV005 (BCRF1, 512 bp) adjacent to the 5’ ITR was highly conserved in all samples. At the 3’ ITR, Clades 2.1 and 2.2 had a 622 bp haplotype at the region homologous to LSDV153 differing from a 555 bp Clade 2.3 haplotype (Figure 3). These divergent 3’ ITR haplotypes’ had a high level of differentiation (similarity of 79%).

**Fig 3.**
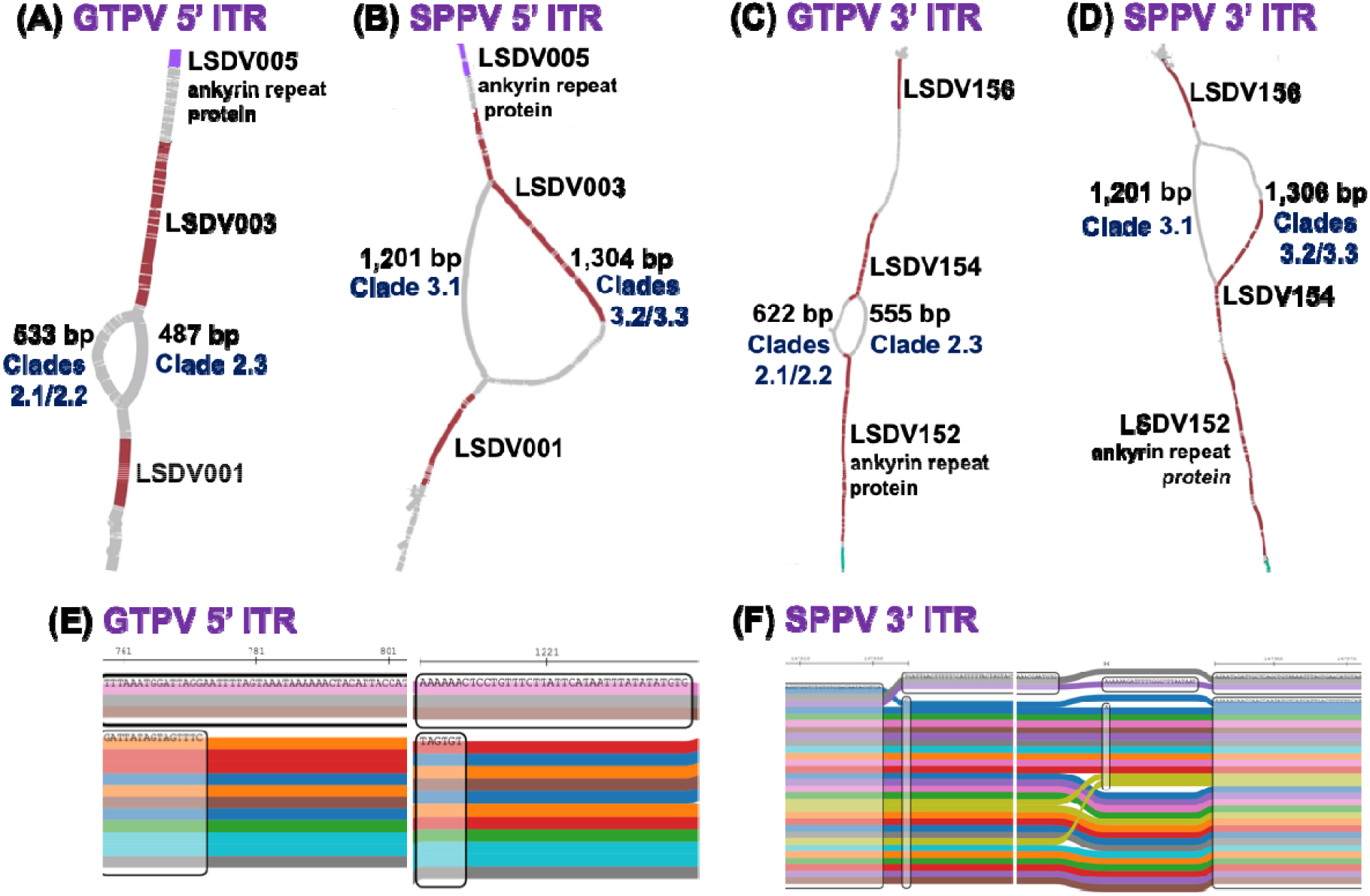
Top: The (A) GTPV 5’ ITR, (B) SPPV 5’ ITR, (C) GTPV 3’ ITR, and (D) SPPV 3’ ITR. (A) The GTPV PVG region homologous to LSDV002 had two divergent haplotypes, separating at node 57 and merging at node 104. One haplotype was 533 bp (in Clades 2.1/2.2) and the other was three nodes spanning 487 bp (in Clade 2.3). (B) The SPPV PVG region homologous to LSDV002 had two divergent haplotypes, separating at node 55 and merging at node 109. One haplotype spanned 1,201 bp (Clade 3.1) at five nodes that had a truncated LSDV003. The other was 48 nodes spanning 1,304 bp and encoded a complete LSDV003 (Clades 3.2/3.3). (C) The GTPV LSDV153 region had two divergent haplotypes: one common to Clades 2.1/2.2 was 622 bp across 50 nodes, and a Clade 2.3 one that was 555 bp across 16 nodes. (D) The SPPV LSDV155 region had two divergent haplotypes. One spanned 1,201 bp (Clade 3.1) across five nodes and had a truncated LSDV154. The other spanned 53 nodes totalling 1,306 bp (Clades 3.2/3.3) with a complete LSDV154. Bottom: SequenceTubeMap images of the PVG ITR regions. (E) The GTPV PVG at 759 to 1,230 bp using the NC_004003 coordinates showing the divergent haplotype in the three Clade 2.3 samples (top three rows). (F) The SPPV PVG at 147,907-972 bp using the NC_004002 coordinates showing the divergent haplotype in the Clade 3.1 samples (top two rows).

The SPPV PVG had two divergent haplotypes at the 5’ ITR: one 1,201 bp in Clade 3.1, and the other 1,304 bp (in Clades 3.2 and 3.3) (Figure 3). These haplotypes were at bases 53-683 of the LSDV003 genes coordinates, and had 86% similarity due to numerous rearrangements, which contrasted with 98-99% similarity for adjacent regions. Vaccine-related Clade 3.1 had a truncated non-functional LSDV003, unlike Clades 3.2 and 3.3. At the 3’ ITR, there were two divergent haplotypes with lengths 1,201 bp (Clade 3.1) and 1,306 bp (Clades 3.2/3.3) in which only the latter encoded a complete LSDV154 (Figure 3). Within SPPV clades 3.2 and 3.3, intra-clade variation produced two alternative putative ORFs: a shorter LSDV003 in 24 samples and all of GTPV and LSDV, and a longer LSDV003 in SPPV six samples with 124 additional amino acids associated with a 46 bp indel (Figure S10). These had identical homology at the 3’ end, implying short and long LSDV154 ORFs in these samples. The longer LSDV003 and LSDV154 isoforms were in four Clade 3.2 (OR239060, AY077832, NC_004002, MN072629) and two Clade 3.3 (PV167793, PV434148) samples.

Evidence for selective processes was evaluated at each gene by comparing the ancestral D_N_ and D_S_ numbers between these two species to those within each species for GTPV and SPPV (P_N_, P_S_) (Stoletzki & Eyre-Walker 2011). At the ITRs, this showed high conservation at LSDV001 and LSDV156, which had no nonsynonymous mutations in any of the GTPV and SPPV samples (Table S3). LSDV001 and LSDV156 promote host IL-1B-related and TNFa-related signalling pathways that facilitate skin nodule formation (Yang et al 2025). The regions similar to LSDV002 and LSDV155 had ancestral conservation as well as no synonymous changes but numerous nonsynonymous ones in GTPV (P_N_/P_S_ 3/0 for each gene) and SPPV (P_N_/P_S_ 13/0). These differences stemmed from the high diversity between the haplotypes at this region. LSDV003 and LSDV154 also had ancestral conservation, and in GTPV there similar levels of nonsynonymous and synonymous change compared to the genome-wide ratio of 0.98. However, in SPPV both had a high rate of nonsynonymous changes (P_N_/P_S_ 10/0 for both) arising from the divergent haplotypes (Figure 3). LSDV004 and LSDV153 encode secreted immunomodulatory virulence factors that are expressed early during infection (Tulman et al 2002): the regions similar to LSDV004 and LSDV153 had high rates of nonsynonymous changes stemming from pseudogenisation (D_N_/D_S_ 10/1 and 26/2, respectively). LSDV004 had moderate conservation within GTPV (P_N_/P_S_ 13/19) and SPPV (P_N_/P_S_ 1/1), as did LSDV153 in SPPV (P_N_/P_S_ 0/0) but not GTPV (P_N_/P_S_ 8/0), which was associated with two divergent haplotypes (Figure 3).

### Genetic variation at genes associated with CaPV species specificity

To explore genetic variation across GTPV and SPPV, diversity at each gene was examined using the LSDV gene nomenclature for consistency, including regions encoding genes in LSDV that are not CDSs in GTPV or SPPV (Figure S11). GTPV genes had more SNPs/Kb than SPPV ones (median 15.3 vs 5.0) and more mutations/Kb/sample (mean 4.66 vs 0.91) (Table S4). The SNPs/Kb (r=0.53) and mutations/Kb (r=0.46) were positively correlated between GTPV and SPPV, particularly at three genes with high levels of diversity in both species: LSDV130, LSDV131 encoding superoxide dismutase-like (SODC) protein that may be a virulence factor (Xie et al 2024), and LSDV152 encoding an ankyrin repeat protein (Figure S12). In SPPV, 18 genes were conserved (0 or 1 SNPs: LSDV023, LSDV031, LSDV040, LSDV046, LSDV052, LSDV055, LSDV060, LSDV067, LSDV072, LSDV078, LSDV092, LSDV097, LSDV104, LSDV106, LSDV111, LSDV113, LSDV120, LSDV126). Only one gene in GTPV had no SNPs (LSDV105), and five others had <5 SNPs/Kb (LSDV044, LSDV093, LSDV095, LSDV096, LSDV107). These could represent genes involved in essential GTPV- and SPPV-specific processes.

A genome-wide scan for selective processes identified additional patterns, including a comparatively low correlation between the GTPV and SPPV DoS metrics (rho=0.23, p=0.0087) and evidence of long-term conservation across genes (median D_N_/D_S_ =0.63). Although the median DoS values were similar (GTPV 0.031 vs SPPV -0.038), the low correlation was driven by a smaller standard deviation (SD) in the DoS values for GTPV compared to SPPV (0.26 vs 0.39). In addition, there was a higher P_N_/P_S_ ratio within SPPV compared to GTPV (0.98 vs 0.56), which could be associated with the relatively more ancient origin of GTPV allowing more time for selection to remove deleterious nonsynonymous alleles. Four genes had DoS > 0.291 in GTPV and DoS > 0.36 in SPPV (corresponding to the respective median+SD values): LSDV004, LSDV023, LSDV068 and LSDV129. In addition to these, five GTPV and 12 SPPV genes met the species DoS criteria (Supplementary Text).

Among the latter 12, LSDV087 and LSDV141 were previously associated with a SPPV Clade 3.3 outbreak in Russia (Krotova et al 2022): LSDV087 encodes a mutF motif protein that decaps mRNAs, and LSDV141 encodes an extracellular enveloped virion (EEV) membrane protein that may support viral formation and spread.

### Genetic variation at non-ITR genes associated with CaPV species specificity

Our PVG-based approach permitted insights into haplotype-level patterns at key genes of interest. We focused on five LSDV non-ITR ORFs that are inactivated in GTPV and SPPV, which have also been associated with moderating host immune responses (LSDV009, LSDV013, LSDV026, LSDV132, LSDV136) (Tulman et al 2002, Wang et al 2025). In GTPV, LSDV009, LSDV013 and LSDV026 has SNP densities in excess of the genes’ median rate + 2*SD (along with LSDV003 and LSDV153). In SPPV, LSDV009, LSDV130 and LSDV132 also had comparably high SNPs/Kb. There were divergent haplotypes at the regions homologous to these five genes differentiating the three GTPV Clades, but none in SPPV. An example was LSDV026: an LSDV-specific gene potentially similar to VACV gene F11L, which affects host cell machinery manipulation and spread (Wang et al 2025). The GTPV LSDV026 region (at 17,410-18,341) had three haplotypes that extended into LSDV025, which encodes a putative Ser-Thr protein kinase based on homology with VACV VPK2 (Figure 4). One haplotype was 565 bp, another was 563 bp and the third was 552 bp, differentiating Clades 2.1, 2.2 and 2.3, respectively (Figure 4). LSDV026 had a high P_N_/P_S_ ratio within GTPV (25/3) and an elevated ancestral D_N_/D_S_ of 5/0, indicating pseudogenisation. In SPPV, the LSDV026 region had two main isoforms that had a large number of stop codons but a normal P_N_/P_S_ ratio (4/3), raising questions about the role of this sequence (Figure S13).

**Fig 4.**
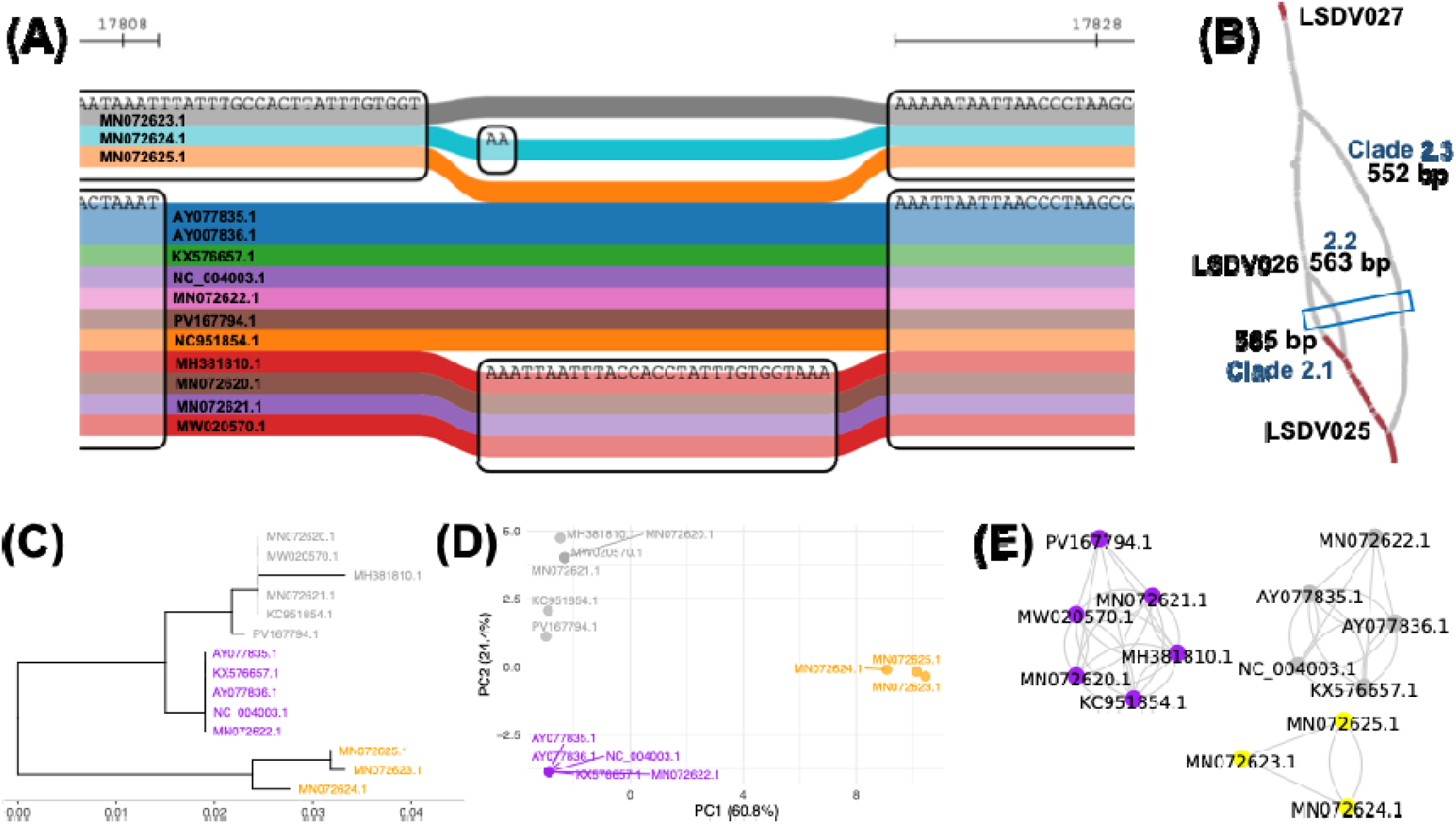
The GTPV PVG region homologous to LSDV026. (A) A sequenceTubeMap image shows Clade 2.3 at the top (MN072623, MN072624, MN072625 in grey, blue and orange), seven Clade 2.1/2.2 samples in the middle, and four samples from Clade 2.2. (B) Bandage graph of three LSDV026 haplotypes of lengths 522 bp, 563 bp and 565 bp with LSDV027 at the top and LSDV025 at the bottom. The blue box indicates the region shown in (A). A full 1,344 bp LSDV025 ORF was present in all genomes. (C) A mid-rooted phylogeny with three clades denoted 2.1 (grey, n=6), 2.2 (purple, n=5) and 2.3 (orange, n=3). Nodes with bootstrap support > 90 are shown. (D) PCA based on SNP data. (E) A network where each node represents one sample. Grey lines indicate inter-community connections, and black lines indicate intra-community ones. The NC_004003.1:17,410-18,341 coordinates for LSDV026 were used.

### Possible signals of convergent evolution in GTPV and SPPV

Recent ancient and pre-modern SPPV genomes (L’Hote et al 2026) further illuminate genetic variation during CaPV evolution and SPPV differentiation through comparison of the shared and unique alleles in the core genomes of LSDV, GTPV and SPPV representative samples. This showed that modern SPPV were the most divergent based on the number of unique alleles, following by GTPV (1,204 vs 1,054 vs 641). In pre-modern SPPV, the same pattern was present, though the number of core alleles shared by all species increased from 95 to 158. The latter metric increased again to 522 when LSDV, GTPV and ancient SPPV were examined. Ancient SPPV (MYRZ180) shared more alleles with LSDV (1,008) than GTPV (544), relative to LSDV and GTPV’s overlap (697). This also showed the relative number of shared SPPV-GTPV alleles increasing from ancient to modern (544 to 564), whereas the level for SPPV-LSDV fell (1,008 to 973), hinting at the potential for the higher genetic similarity of sheep and goat host immune systems to shape CaPV evolution convergently and differently from that of cattle.

In the coding L’Hote et al (2026) SPPV data, there were 627 mutations where the ancient SPPV allele matched LSDV and GTPV, but differed with pre-modern and modern SPPV, and also 245 CDS mutations where the ancient and pre-modern SPPV allele matched LSDV and GTPV, but differed from modern SPPV (Table S5). 327 amino acid alleles where both pre-modern and modern SPPV were identical and differed from ancient SPPV (excluding pseudogenes) were more often identical to GTPV than to LSDV (16 vs 7). Similarly, of the 110 amino acid alleles in modern SPPV differing from ancient and pre-modern SPPV (excluding pseudogenes), more were like GTPV than LSDV (9 vs 1). This implicated 33 amino acid substitutions in possible convergent adaptations (Table S6). Notably, two genes had changes in both GTPV-related subsets: both had two amino acid alterations in the ancient to pre-modern transition, followed by another in the pre-modern to modern period. These genes were: LSDV032 encoding a poly(A) polymerase large subunit, and LSDV042 encoding a serine/threonine kinase (Tulman et al 2022). Collectively, these results underpinned the high differentiation and putative convergent evolution of SPPV and GTPV.

## Discussion

This study explored the evolution and genomic diversity of GTPV and SPPV using phylogenetics, PVG-based and gene-specific approaches. GTPV and SPPV cause extensive disease outbreaks, along with LSDV (WOAH 2010). Understanding better the genetic basis for these differences is pivotal. This study revealed clear contrasts in population structure, evolutionary history, and genome evolution between GTPV and SPPV. GTPV had a deeper and more stable population structure, comprising three genetically distinct clades with little evidence of recent admixture, whereas SPPV was a mosaic of more recent variation with weaker clade separation. For both viruses, genome-wide diversity patterns were consistent with an excess of rare mutations.

This work builds on earlier gene-centric CaPV pangenome analyses (Xie et al 2024) by using PVGs that did not rely on gene annotation, capture non-coding and structural variation, and quantified PVG openness using multiple complementary approaches. Importantly, pangenome openness here reflected both biological diversity and uneven sampling, particularly the limited representation of samples from Africa, which was a constraint for both species. Across all genomes, the shared number of genomic bases was smaller for GTPV than for SPPV (142.4 vs 144.6 kb), reflecting GTPV’s stronger geographic structure. However, diversity at the 5′ and 3′ genome ends was higher relative to the core genome in SPPV than GTPV, consistent with the 5’ and 3’ end genes’ roles in encoding non-essential virulence and host immunomodulatory factors (Tulman et al 2002). The open PVG inferred for SPPV indicated that broader geographic and temporal sampling will continue to reveal novel variation, whereas the closed GTPV PVG suggested limited returns from additional sampling in Eurasia. Lastly, the representative PVGs constructed here can be a practical resource for improved short-read mapping and variant detection in future CaPV work (Wright et al 2025).

PVG-based analyses were particularly informative at the ITRs, revealing substantial structural and haplotypic variation centred on regions homologous to LSDV003 and LSDV154. These genes are annotated as encode ER-localised apoptosis regulators that interfere with host mitochondrial function and apoptosis (Tulman et al 2001). LSDV003 is predicted to have a signal peptide, a transmembrane domain, and insert into the endoplasmic reticulum (Liaci & Förster 2021). The protein it encodes is similar to MYXV M004, an essential virulence factor (Enow 2025). The LSDV003 and LSDV154 truncations in vaccine-related SPPV Clade 3.1 were consistent with attenuated virulence following serial passage in non-natural hosts. Variation at LSDV154 was previously associated with SPPV outbreaks in south-western Europe (Breman et al 2024). The longer putative LSDV003 ORFs in six SPPV samples across Clades 3.2 and 3.3 may be associated with expression regulation of this protein via protease cleavage. This potential structural ITR plasticity could mean that these extended ORFs encode mRNAs with different expression levels or post-translational functions. In addition, the retention of the non-CDS regions suggests a continued functional relevance, perhaps to facilitate recombination for those at the ITRs.

Beyond the ITRs, patterns of gene-specific diversity highlighted candidate loci associated with host specificity. 18 SPPV genes had either zero or one SNP, implying strong functional constraint. In contrast, GTPV only had one gene with high conservation, suggesting selective processes maybe operating on each clade differently, perhaps linked to their higher geographic population structure. There were signals consistent with adaptive evolution at three genes, including LSDV068 encoding a polyA polymerase small subunit. The second was: LSDV023, which had sequence-based and structural homology on viro3D (Litvin et al 2025) with MYXV *m018L* gene that encodes non-essential cytoplasmic protein (Higley & Way 1997). The third was LSDV129, which had similarity to MYXV M130R, whose product has substantive immunomodulation and virulence effects in rabbit hosts (Barrett et al 2009). In addition, the three major GTPV lineages were consistently differentiated by divergent haplotypes at regions corresponding to LSDV002, LSDV009, LSDV013, LSDV026 and LSDV153. In addition, this study found possible evidence of convergent evolution in SPPV and GTPV using published ancient and pre-modern SPPV genomes (L’Hote et al 2026). This implied that the higher similarity of the goat and sheep immune systems relative to that of cattle since the Bronze Age may elicit shared changes at specific proteins, their genes and even at the nucleotide level. Here, 16 and nine such SPPV amino acid substitutions became GTPV-like in the Bronze Age to pre-modern and pre-modern to modern eras, respectively, whereas comparable numbers for LSDV were seven and one, respectively. One of the proteins implicated (at LSDV032) is a polyA polymerase component, like LSDV068’s gene product above.

In summary, GTPV evolution is structured by deep, clade-specific divergence, whereas SPPV has a more diffuse ongoing diversification from a more recent origin. Several broader questions emerge from this study. First, the retention of long non-coding regions at loci that are genes in LSDV only (LSDV002, LSDV004, LSDV009, LSDV013, LSDV026, LSDV132, LSDV153, LSDV155) suggested that these regions may be subject to functional or structural constraints beyond protein coding. ITRs are constrained by replication mechanisms, whereas the preservation of internal non-CDS loci across divergent lineages argues for targeted functional testing. Second, many of the complex rearrangements described here, especially at the ITRs, are likely under-resolved in short-read assemblies. Long-read sequencing and haplotype-resolved assemblies will be essential for improving CaPV genome annotation, resolving ORF structure, and linking genomic variation to viral phenotype. Third, there were vaccine-linked specimens with unique ITR mutations: these recombination-prone regions may be as hotspots for new adaptive variation, underscoring the need for genomic surveillance of vaccine-linked strains using PVGs (Guarracino et al 2023). Fourth, GTPV isolates infect both sheep and goats (Biswas et al 2019) and yet SPPV may be the most basal among the CaPV (L’Hote et al 2026), highlighting continued evolutionary questions that may impact perspectives on host tropism.

## Supplementary Material

Supplementary Table 1. The genomes examined showing their species, clade, accession, genome length, number of annotated CDSs, sample namer, year of isolation, host of isolation, country of isolation and context (field vs vaccine).

Supplementary Table 2. The matching gene coordinates for GTPV and SPPV based on LSDV. The gene ID for each species is shown, along with the gene starts and ends. Pseudogenes in GTPV and SPPV that are CDSs in LSDV are denoted as “inferred”.

Supplementary Table 3. Direction of selection metrics for each gene, showing the strand, number of ancestral nonsynonymous changes (dN), number of ancestral synonymous changes (dS), number of population-level nonsynonymous changes in GTPV (pN_G), number of population-level synonymous changes in GTPV (pS_G), direction of selection value in GTPV (DoS_G), number of population-level nonsynonymous changes in SPPV (pN_S), number of population-level synonymous changes in SPPV (pS_S), and direction of selection value in SPPV (DoS_S).

Supplementary Table 4. The numbers of GTPV SNPs/Kb, GTPV mutations/Kb, SPPV SNPs/Kb and SPPV mutations/Kb.

Supplementary Table 5. The allelic states at variable sites among LSDV (NI_2490 AF325528, isolated in 1958), GTPV (NC_004003, isolated in 2000), modern SPPV (MW167071, isolated in 2018), pre-modern SPPV (CL299) and ancient SPPV (MYRZ180).

Supplementary Table 6. 33 amino acid changes associated with convergent adaptations. The table outlines the site in LSDV NI_2490 AF325528 (Site_LSDV), the context (time period of ancient to pre-modern or pre-modern to modern; and species GTPV or LSDV), the LSDV, GTPV, ancient SPPV, pre-modern SPPV and modern SPPV nucleotide alleles, the gene name, mutation type, gene nucleotide base and amino acid site, the LSDV, GTPV, ancient SPPV, pre-modern SPPV and modern SPPV amino acid alleles, and the LSDV, GTPV, ancient SPPV, pre-modern SPPV and modern SPPV codons.

Supplementary Table 7. The primers examined, showing the virus or subset, primer orientation, primer name, primer sequence, start in LSDV, end in LSDV, LSDV gene region, LSDV gene product, number of LSDV mismatches, start in SPPV NC_004002, end in SPPV NC_004002, SPPV NC_004002 gene region, SPPV NC_004002 gene product, number of SPPV NC_004002 mismatches, start in GTPV NC_004003, end in GTPV NC_004003, GTPV NC_004003 gene region, GTPV NC_004003 gene product, and number of GTPV NC_004003 mismatches.

Supplementary Table 8. Selected genes of interest in this study. The gene LSDV identifier, gene name, reason for inclusion (Purpose), Vaccinia virus homolog (VACV) and synonyms.

## Acknowledgements

This study acknowledges the use of resources from Pirbright Institute’s Bioinformatics STP and the Dept of Biology at the University of the Philippines Baguio, and thanks Tiffany Tobi, Mai Prihartini, Jonas Albarnaz, Miguel Hernandez and Caroline Wright for useful discussions.

## Data Availability

All supporting code and protocols have been provided within the article, through supplementary data files is available at https://github.com/downingtim/GTPV-SPPV-PVG

