## Supplementary_data for "Insights into goatpox virus and sheeppox virus genomes from pangenome graphs"

**Section	Page**

Supplementary Results	2

Figure S1	5

Figure S2	6

Figure S3	7

Figure S4	8

Figure S5	9

Figure S6	10

Figure S7	11

Figure S8	12

Figure S9	13

Figure S10	14

Figure S11	15

Figure S12	16

Figure S13	17

Figure S14	18

Figure S15	19

Figure S16	20

Figure S17	21

Figure S18	22

**Supplementary Results**

**Genetic variation at genes associated with attenuated virulence**

Genes associated with attenuated virulence were conserved and retained low or moderate levels of variability. These genes encoded ankyrin-repeat proteins (LSVD012, LSVD145, LSVD147, LSVD148); a kelch-like proteins (LSDV019, LSDV144, LSDV151), a B22R-like protein (LSDV134) and host range factors (LSDV067, LSDV141). LSDV134 is a 6.1 Kb gene encoding a variola virus B22R-like protein similar to mpox protein MPXV197 that inhibits host immune responses via T-cell maturation (Alzhanova et al 2024). In Clade 3.1, this was fragmented into putative CDSs of 2.4, 0.2, 0.5, 0.4 and 2.4 Kb, suggesting an absence of a functional B22R protein, which is associated with vaccine-related CaPV (Berguido et al 2023) and attenuated virulence (Alzhanova et al 2024).

**Genetic variation at genes associated with CaPV species specificity - LSDV009, LSDV013, LSDV132 & LSDV136**

LSDV009 encodes putative alpha amanitin-sensitive protein from the A52R-like family in LSDV that has homology with the N2L protein. There were three distinct haplotypes at LSDV009 (at NC_004003.1:5,583-5,899): one (359 bp) unique to Clade 2.3, another (271 bp) in five samples (AY077835.1, AY077836.1, KX576657.1, MN072622.1, NC_004003.1), the third (522 bp) in the other five Clade 2.1 samples, and PV167794.1 had a mix of these haplotypes (Figure S14).

LSDV013 encodes an interleukin-1 receptor (IL-1R) homolog, which can bind and inactivate host IL-1β, but is truncated some LSDV, as well as GTPV and SPPV (REF). In GTPV, the region similar to this gene (at NC_004003.1:8,395-9,465) had a high rate of SNPs with two haplotypes: one 735 bp in Clade 2.3 and the other 721 bp in Clades 2.1 and 2.2 (Figure S15). Within GTPV and SPPV, LSDV013 had a high rate of nonsynonymous changes, symptomatic of pseudogenisation.

The region similar to GTPV LSDV132 had a high rate of nonsynonymous SNPs (PN/PS 3/0) and structural changes with limited clade-related structure (Figure S16). SPPV LSDV132 had high rate of nonsynonymous SNPs too (PN/PS 16/0), as did the ancestral DN/DS rate (17/0). This low rate of conservation was consistent with pseudogenisation, like previous work (Xie et al 2024). In LSDV, LSDV132 supports viral replication by delayed host cell apoptosis (Wang et al 2025).

GTPV LSDV136 had a high level of population structure, suggestive of long-term conservation within clades, and was likely had a functional ORF in Clades 2.1 and 2.2 (Figure S17). In SPPV, most diversity at LSDV136 was within Clade 3.1 (Figure S13). Diversity within LSDV136 was conserved within GTPV (PN/PS 6/6) but not SPPV (PN/PS 11/0), consistent with continued function in GTPV but not in SPPV.

**Genetic variation at genes associated with diagnostics**

Diversity at three regions associated with diagnostic assay differentiating CaPV were examined (Table S7). The first was LSDV074, a p32 gene encoding an IMV envelope protein that is a VACV H3L homolog. The second was LSDV036, a RPO30 gene encoding an RNA polymerase 30 kDa subunit that is a VACV E4L homolog with a 21 bp deletion unique to SPPV (Lamien et al 2011). The third was LSDV146, a G-protein-coupled chemokine receptor aka GPCR gene encoding a phospholipase-D-like protein.

In addition, diversity at regions specific to either SPPV (LSDV045 encoding a DNA-binding phosphoprotein similar to VACV I3L), GTPV (LSDV099 encoding an intermediate transcription factor VITF-3), the WT LSDV Clade 1.2 (LSDV126 encoding an EEV glycoprotein similar to VACV A36R; and LSDV133 encoding a DNA ligase-like protein similar to VACV B22R), and vaccine-linked LSDV Clade 1.1 (LSDV144 encoding a kelch-like protein) were explored. Known gene synonyms are in Table S8.

**Gene-specific patterns of selection**

Five genes were under ancestral positive selection that were conserved in GTPV but not SPPV: LSDV069 encoding a RNA polymerase subunit essential for viral replication (Chen et al 2025); LSDV070 encoding another IMV membrane protein potentially involved in cell entry; LSDV074 encoding an intracellular mature virion (IMV) envelope protein essential for virus morphogenesis; LSDV088 encoding a NTPase/ATPase (NPH-I) essential for virus survival; LSDV136 encoding a protein in the A52R family that may inhibit apoptosis or pro-inflammatory transcription factor activation (Tulman et al 2002).

Twelve genes had signals of ancestral adaptive evolution that were conserved in SPPV but not GTPV: LSDV005 encoding an IL-10-like protein that mimics IL-10 to downregulate host immune responses; LSDV056 encoding a VACV M51R homolog; LSDV064 encoding a membrane protein required for cell entry; nearby LSDV067 encoding a host range protein that moderates host immune responses; LSDV087 encoding a mutT motif protein that decaps mRNA; LSDV095 encoding a virion core protein essential for virus morphogenesis; LSDV104 encoding another IMV envelope protein; LSDV112 encoding a DNA polymerase processivity factor involved in DNA replication; LSDV128 encoding a CD47-like protein that likely mimics host CD47 to moderate associated immune responses (Tulman 2002); LSDV130 encoding a hypothetical protein that may be a calcium-binding regulatory protein; LSDV141 encoding an extracellular enveloped virion (EEV) membrane protein that may have a role in virus formation and spread; and LSDV142 encoding a protein with a Bcl-2 domain that inhibits type 1 INFB by binding IRF3.

LSDV039 encodes a high-fidelity DNA polymerase, which is essential for replication and recombination (Moss 2024). This gene had one of the highest rates of nonsynonymous changes in GTPV, LSDV and SPPV, and yet was conserved within GTPV (PN/PS 69/122) and SPPV (PN/PS 25/26) and on the ancestral lineage (20/54) (Figure S16). At this gene, GTPV was the most ancient and had the most diversity.

**Gene-specific patterns of adaptation in LSDV067 (C7L)**

SPPV had LSDV067 (C7L) changes that were likely to have limited functional effects: S23 (TCC) compared to A23 (GCC) in GTPV (and LSDV), along with I27 (ATT) instead of V27 (GTT) in GTPV (and LSDV). In GTPV compared to SPPV (and LSDV), it has N134 (AAC) versus D134 (GAC), V181 (GTG) versus A181 (GCG), and V192 (GTG) versus C192 (TGC). V192 potentially has a substantial effect. GTPV and SPPV share A210 (GCC) compared to T210 (ACC) in LSDV, along with lacking a glycine-rich expansion at LSDV amino acids 285+ that may have a large effect.

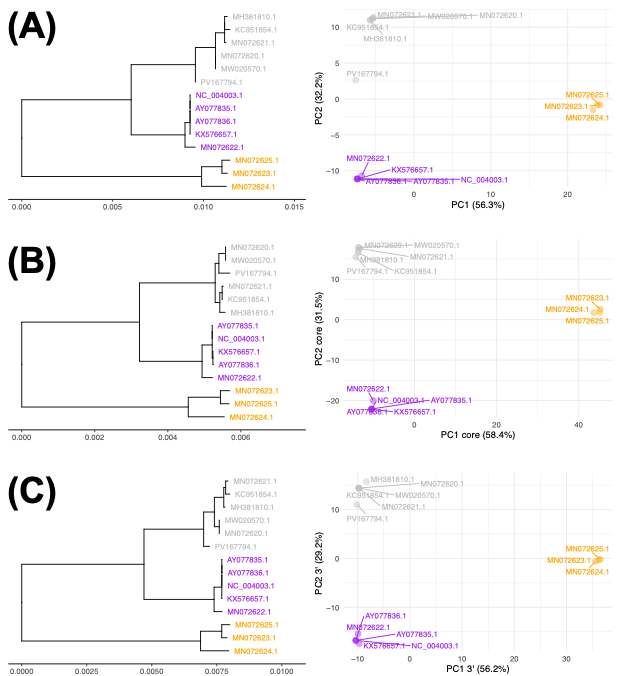

**Figure S1.** Midpoint-rooted phylogenies (left) of the GTPV (A) core genome (365 SNPs), (B) 5’ end (262 SNPs) and (C) 3’ end (392 SNPs) showing three clades: 32.1 (grey, n=6), 2.2 (purple, n=5) and 2.3 (orange, n=3). The scale bar shows the numbers of substitutions per site. Right: a PCAs (right) where PC1 had 56-58% of variation, and PC2 had 29-32% of it.

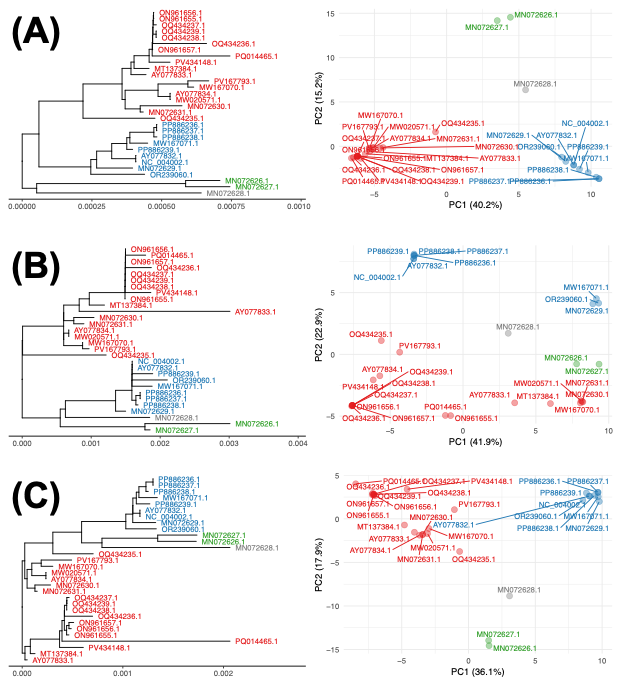

**Figure S2.** Midpoint-rooted phylogenies (left) of the SPPV (A) core genome (720 SNPs), (B) 5’ end (254 SNPs) and (C) 3’ end (341 SNPs) showing three clades: 3.1 (green, n=2), 3.2 (blue, n=9), 3.3 (red, n=18), and MN072628.1 (grey). The scale bar shows the numbers of substitutions per site. Right: a PCAs (right) where PC1 had 36-42% of variation, and PC2 had 15-23% of it.

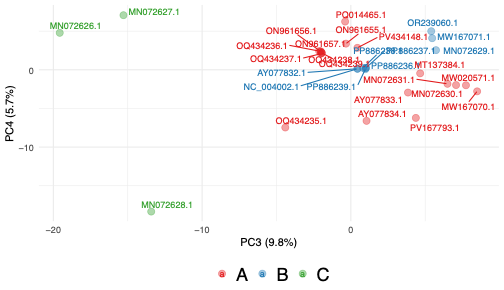

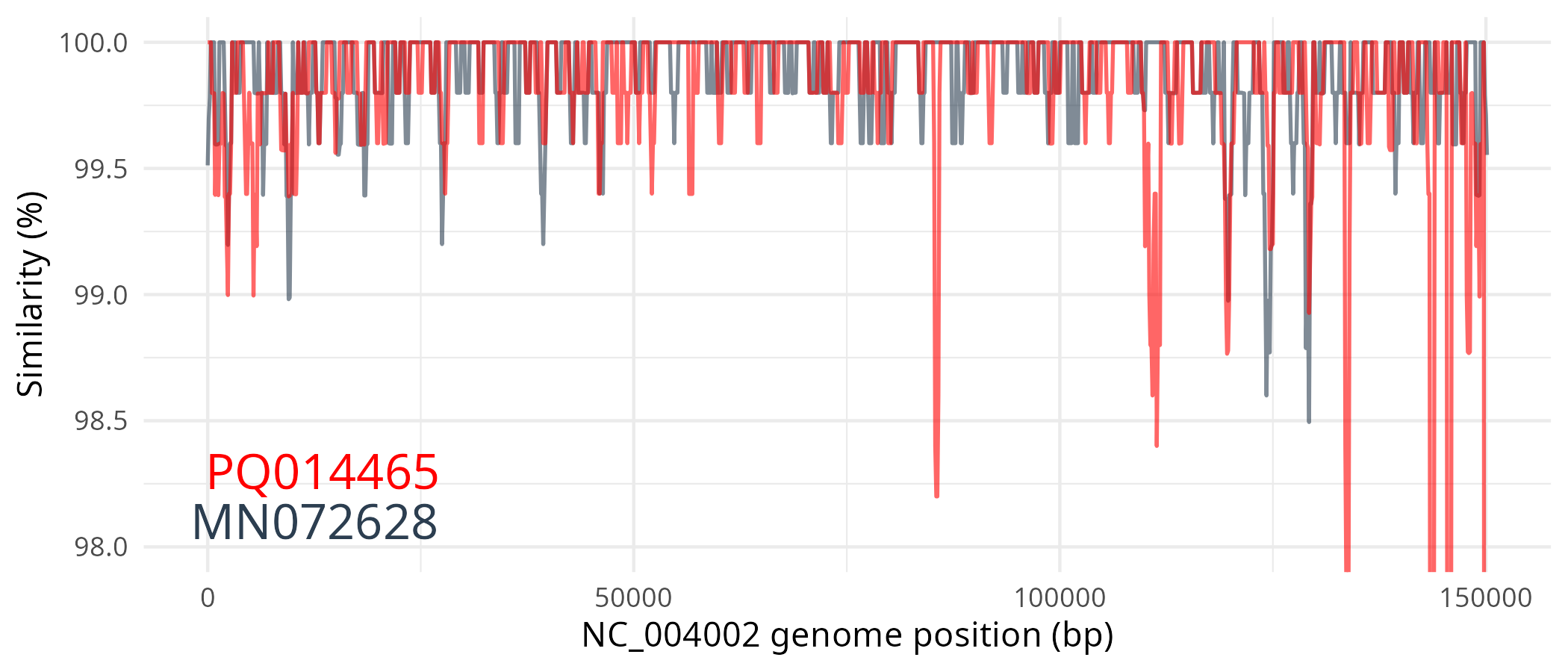

**Figure S3**. Top: PCA of PC3 vs PC4 based on genome-wide SNPs in SPPV (n=30 samples), where PC3 had 10% of variation, and PC2 had 6% of it. The 3 clades were 3.3 (red, n=18), 3.2 (blue, n=9) and 3.1 (green, n=2). Bottom: The relative genome-wide divergences of MN072628 (grey) and PQ014465 (red) compared to NC_004002.

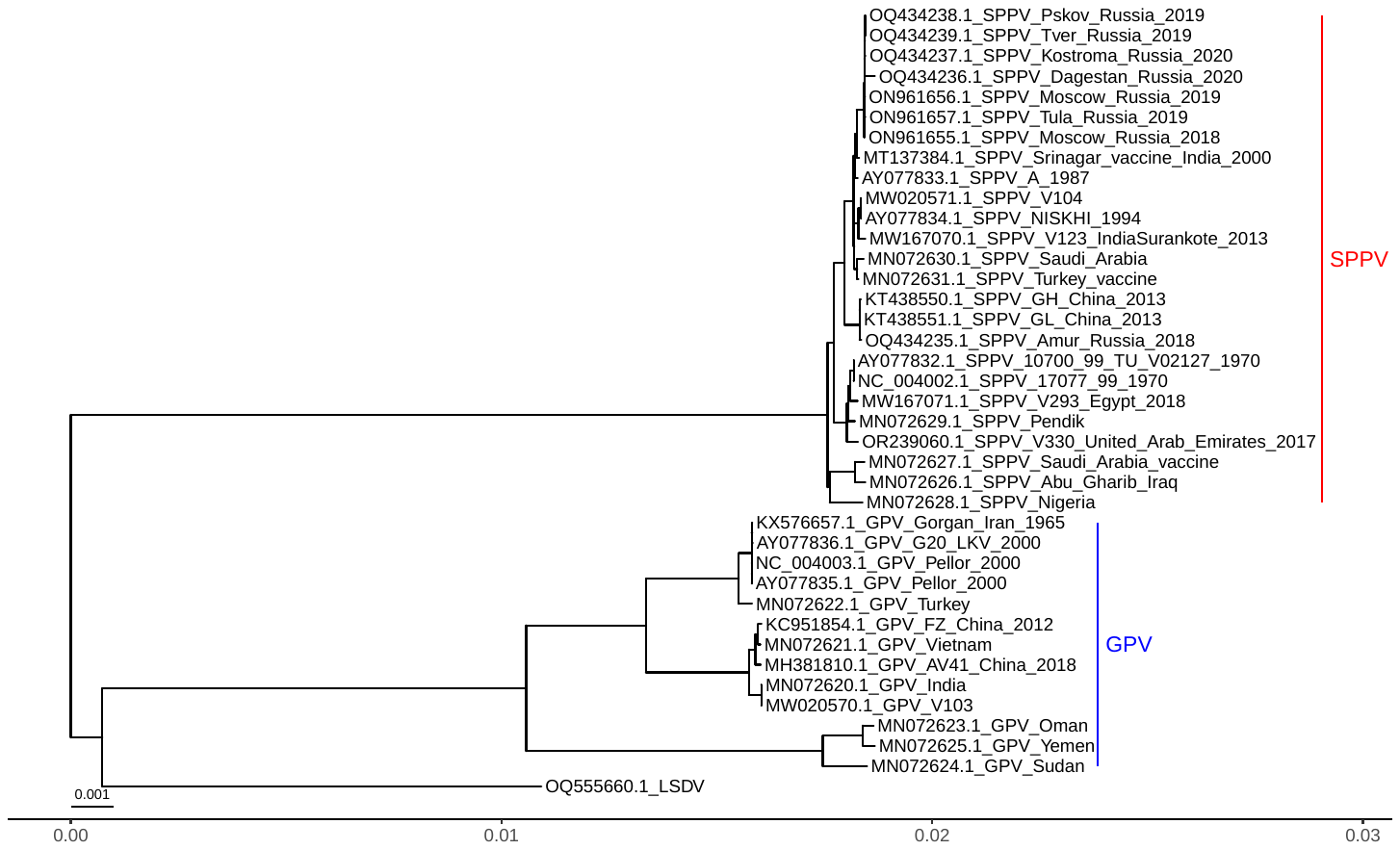

**Figure S4**. A phylogeny based on genome-wide SNPs of valid SPPV (n=25, red) and GTPV (n=13, blue). GTPV was marginally more closely related to LSDV represented by one sample (OQ555660.1). The scale bar shows the numbers of substitutions per site.

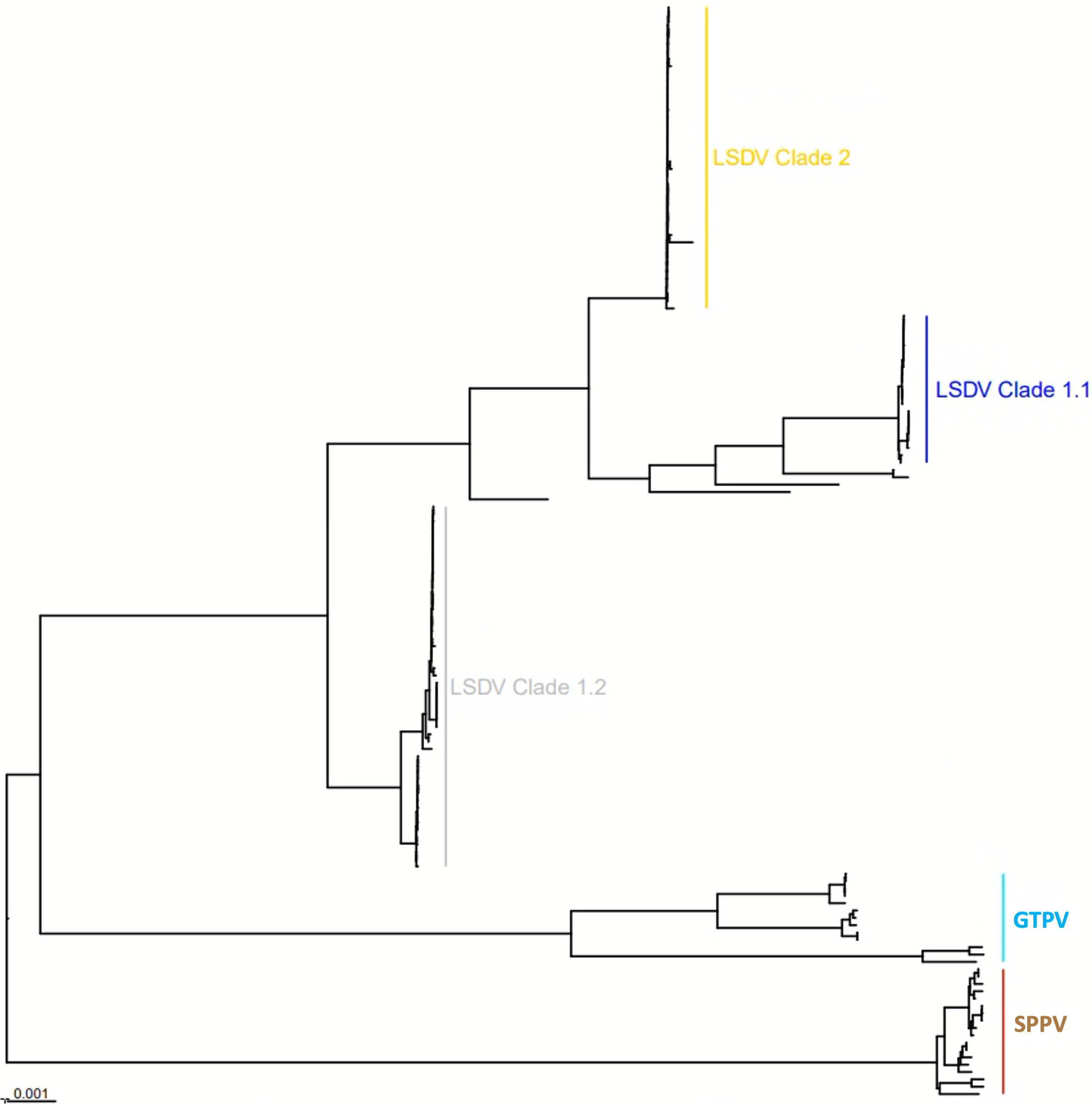

**Figure S5**. A phylogeny based on genome-wide SNPs of n=154 CPPV samples. GTPV (n=13, cyan) was marginally more closely related to LSDV than SPPV (n=25, brown). LSDV is divided into three clades: 1.2 (grey), 1.1 (blue) and 2 (gold). The scale bar shows the numbers of substitutions per site.

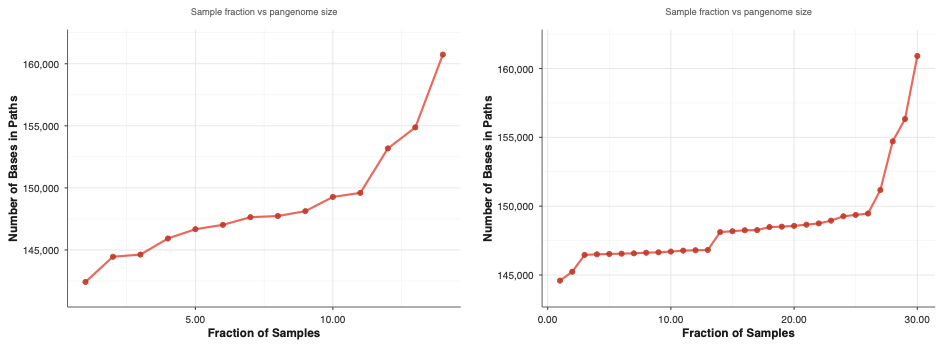

**Figure S6**. The relationship between the average number of samples in the PVG (x-axis) and the number of bases in the corresponding set of paths (y-axis) for GTPV (left) and SPPV (right) using ODGI heaps. The shared PVG (paths found in all samples) was 142,411 bp for GTPV and 144,593 bp for SPPV. The median PVG length was 147,684 bp for GPTV and 148,183 bp for SPPV. The complete PVG (paths found in any sample) was 160,732 bp for GTPV and 161,135 bp for SPPV. The y-axes ranges for these plots differ.

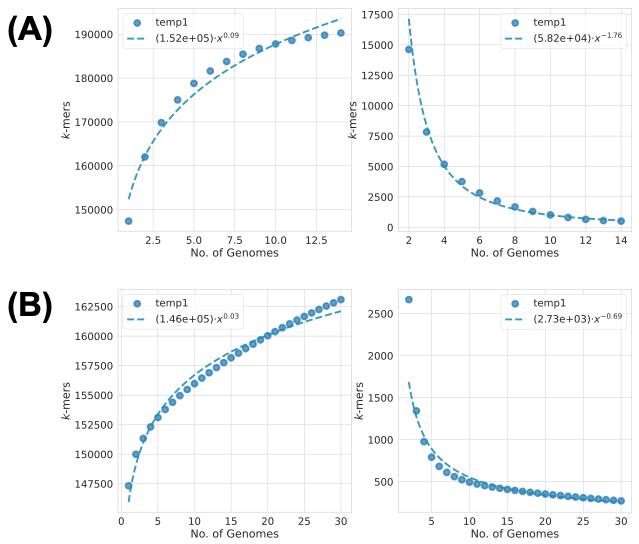

**Figure S7**. PVG growth in terms of the effect of adding more samples (x-axes) on the numbers of new k-mers in the PVG (y-axes) using Pangrowth for (A) GTPV and (B) SPPV. A new k-mer represents a new mutation. This found evidence of a closed PVG for GTPV: gamma=0.09 and alpha=1.76, and an open PVG for SPPV: gamma=0.03 and alpha=0.69 for SPPV.

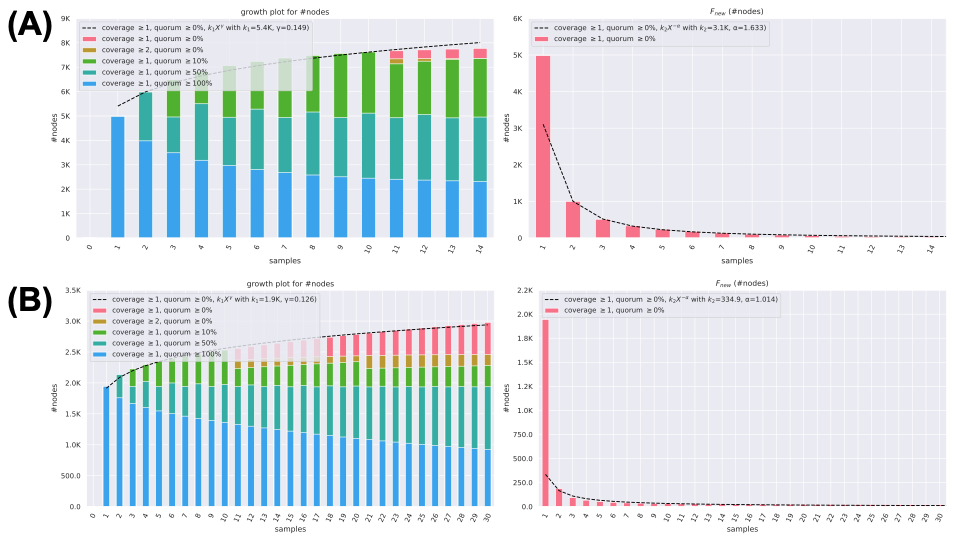

**Figure S8**. PVG growth in terms of the effect of adding more samples (x-axes) on the numbers of (A) total and (B) new nodes in the PVGs for (A) GTPV and (B) SPPV using Panacus. This found evidence of a closed PVG for GTPV (gamma=0.15 and alpha=1.63). This suggested the rate of gain of new mutations was zero when a substantial number of genomes were collected. For SPPV, the PVG was between closed and open (gamma=0.13 and alpha=1.01), suggesting it may depend on the lineage sampled to determine the rate of mutation gain as the number of samples increases.

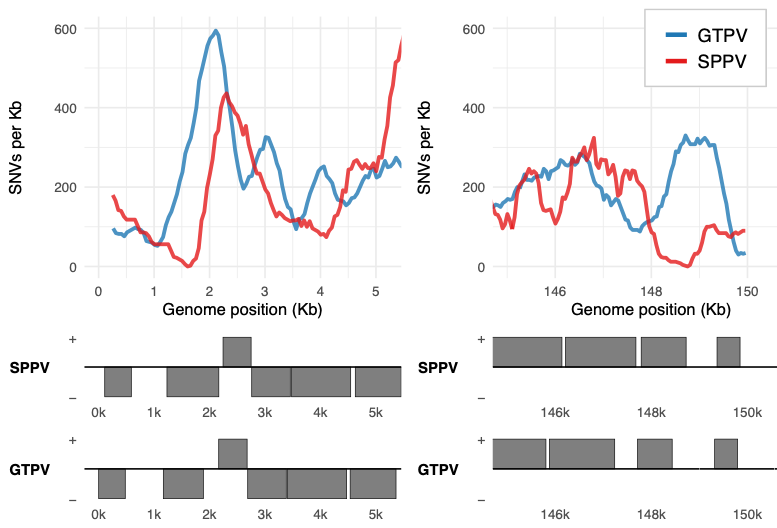

**Figure S9**. The genome-wide SNP density (SNPs/Kb) in GTPV (blue) and SPPV (red) for the 1^st^ and last 5 Kb regions of the genomes. Annotation was derived from the NC_004002.1 and NC_004003.1 genomes. GTPV and SPPV had higher diversity at LSDV_03 and LSDV005 (an ankyrin repeat protein) at the 5’ ITR, and LSDV154 and LSDV156 at the 3’ ITR.

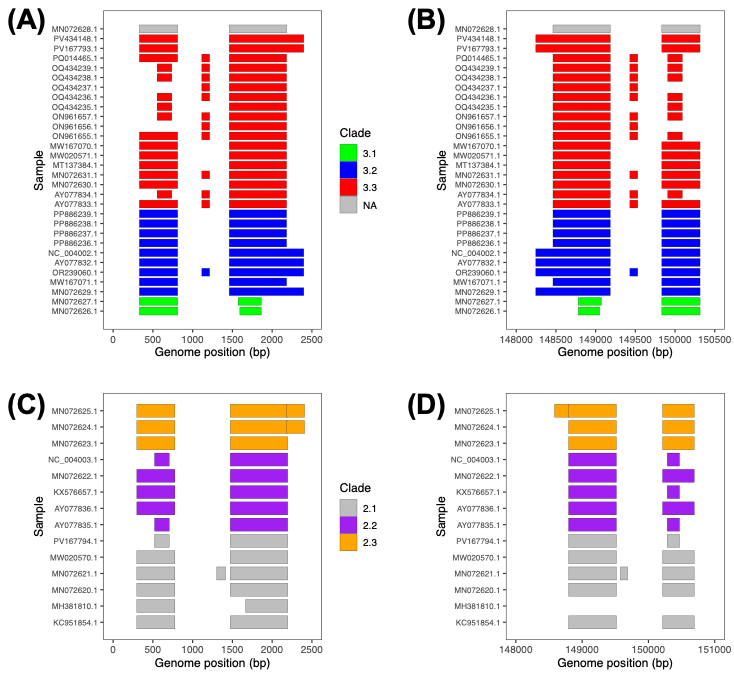

**Figure S10.** Predicted CDSs at the SPPV 5’ (A) and 3’ (B) ITRs, along with the GTPV 5’ (C) and 3’ (D) ITRs. The colours correspond to the clades: SPPV - 3.1 in green, 3.2 in blue, 3.3 in red, NA in grey; and GTPV - 2.1 in grey, 2.2 in purple, 2.3 in orange. The CDS predictions were from Prokka. LSDV003-LSDV004 was a single ORF in six SPPV and two GTPV isolates, with the same for LSDV153-LSDV154. LSDV003 and LSDV154 are truncated in SPPV Clade 3.1. No sample had functional LSDV002 or LSDV155 genes. MH381801.1 (Clade 2.1) lacked a proper 3’ ITR, likely due to assembly issues.

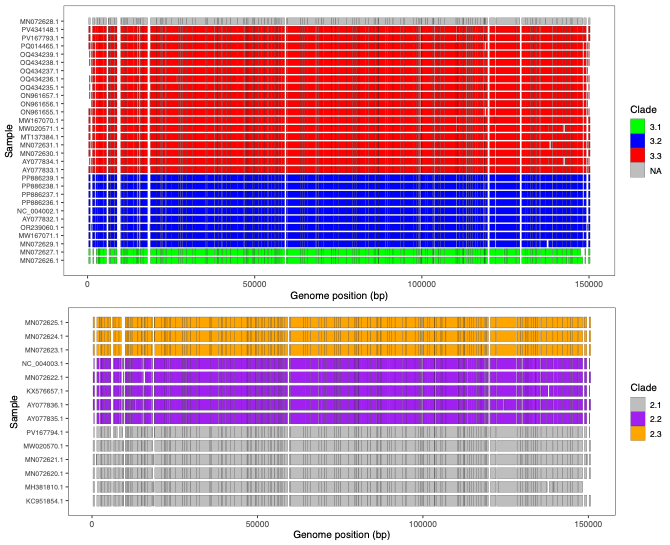

**Figure S11**. Putative CDSs in (A) SPPV and (B) GTPV where the colours correspond to the clades (GTPV: 2.1 in grey, 2.2 in purple, 2.3 in orange) (SPPV: 3.1 in green, 3.2 in blue, 3.3 in red, NA in grey). The CDSs are based on Prokka predictions.

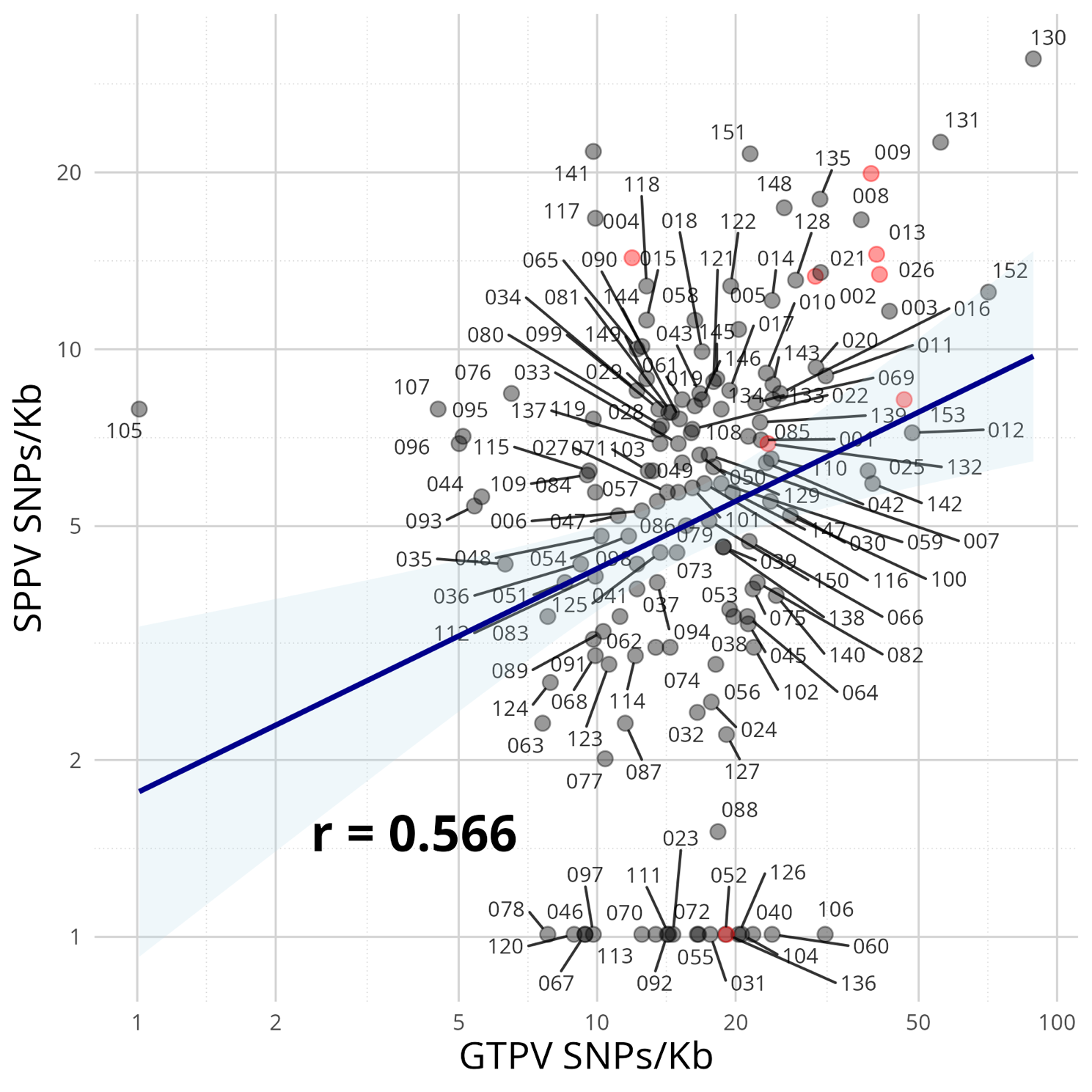

**Figure S12**. The GTPV and SPPV SNPs/Kb was positively correlated (r=0.53) but with wide variation. In SPPV, 18 genes had no SNPs (LSDV023, LSDV031, LSDV040, LSDV046, LSDV052, LSDV055, LSDV060, LSDV067, LSDV072, LSDV078, LSDV092, LSDV097, LSDV104, LSDV106, LSDV111, LSDV113, LSDV120, LSDV126) along LSDV136, which was annotated as a putative pseudogene. One gene in GTPV had no SNPs (LD105), and two had <5 SNPs/Kb (LSDV096 and LSDV107). LSDV130, LSDV131 and LSDV152 had high SNP densities in both species. LSDV002, LSDV004, LSDV009, LSDV013, LSDV026, LSDV132, LSDV136, LSDV153 and LSDV155 are highlighted by red nodes.

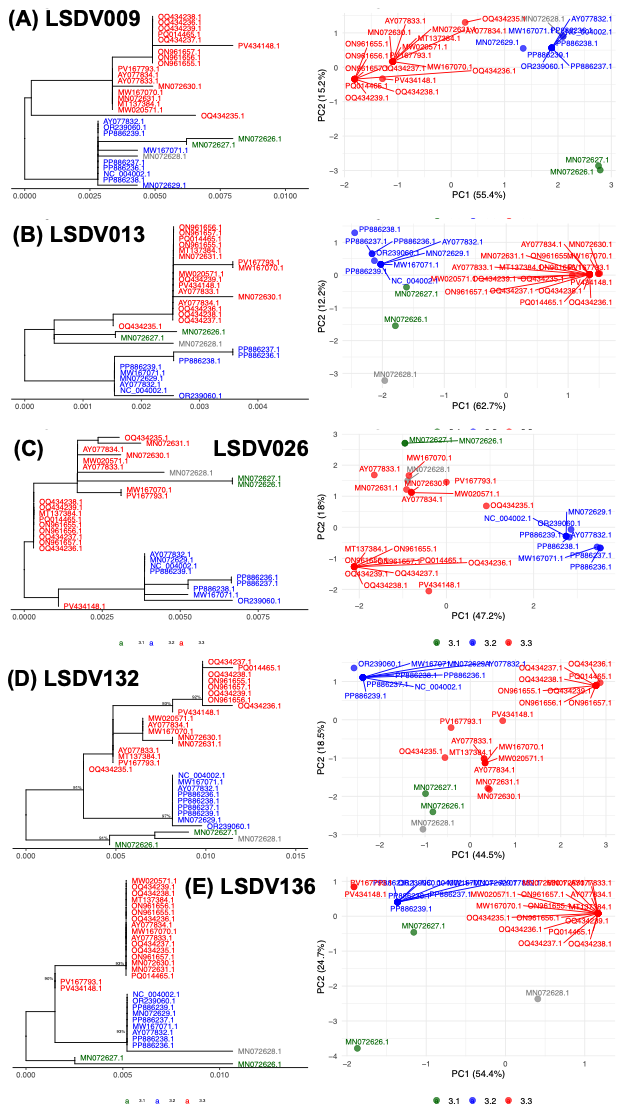

**Figure S13**. SPPV diversity at (A) LSDV009, (B) LSDV013, (C) LSDV026, (D) LSDV132 and (E) LSDV136 showing (left) midpoint-rooted SPPV phylogenies with three clades (3.1 in green, n=2; 3.2 in blue, n=9, 3.3 in red, n=18) and (right) PCA visualisations where the fraction of variation explained is shown on the axes.

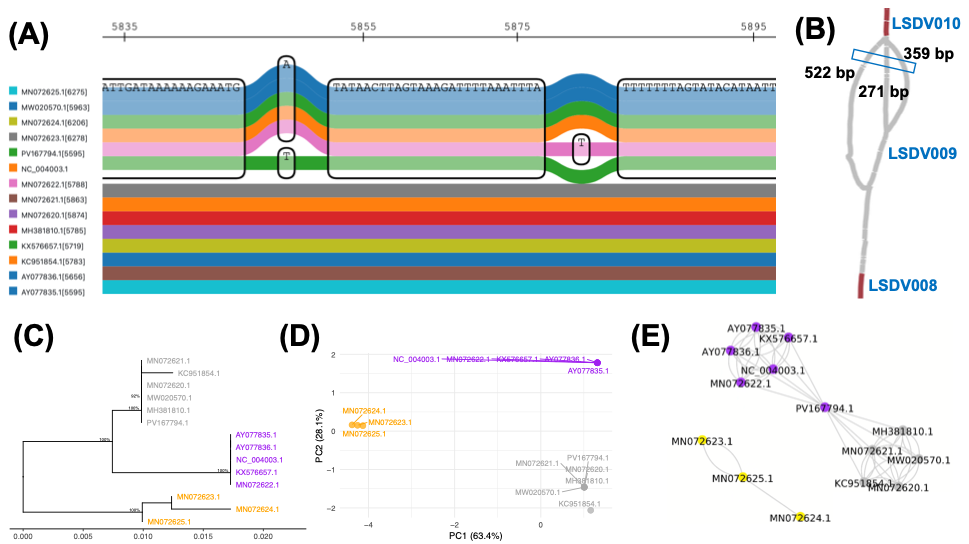

**Figure S14**. Visualisations of the region similar to LSDV009 (at NC_004003.1:5,583-5,899) gene in the GTPV PVG with three haplotypes of lengths 271 bp (in five Clade 2.2 samples, along with PV167794.1 for certain segments), 359 bp (in the three Clade 2.3 samples) and 522 bp (in five Clade 2.1 samples). (A) A sequenceTubeMap image shows an excerpt from the PVG at this region where five samples from Clade 2.2 (AY077835.1, AY077836.1, KX576657.1, MN072622.1, NC_004003.1) are at the top (blue, blue, green, orange, mauve), along with PV167794.1 (green), and the other genomes at the bottom. (B) Bandage image of LSDV009 with LSDV010 is at the top and LSDV008 is at the bottom. The blue box indicates the region shown in (A). (C) A mid-rooted phylogeny with three clades denoted 2.1 (grey, n=6), 2.2 (purple, n=5) and 2.3 (orange, n=3). Nodes with bootstrap support > 90 are shown. (D) PCA based on SNP data. (E) A network where each node represents one sample. Grey lines indicate inter-community connections, and black lines indicate intra-community ones.

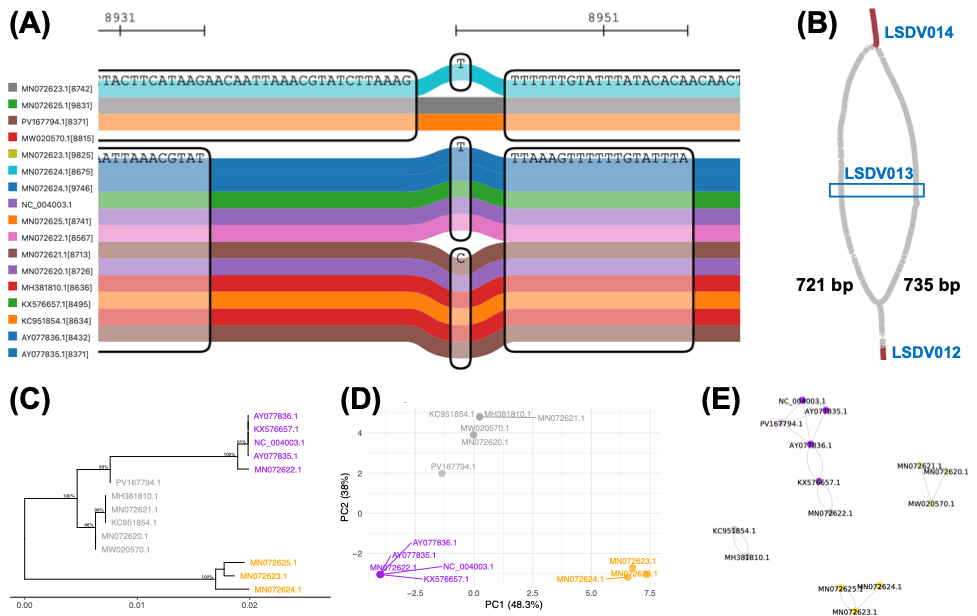

**Figure S15**. Visualisations of the region similar to LSDV013 (at NC_004003.1:1:8,395-9,465) gene in the GTPV PVG with two haplotypes of lengths 721 and 735 bp. (A) The sequenceTubeMap image shows an excerpt from the PVG at this region where MN072623, MN072624, MN072625 at the top (cyan, grey, orange) and the other 11 genomes at the bottom. (B) Bandage image of LSDV013 with LSDV014 is at the top and LSDV012 is at the bottom. The blue box indicates the region shown in (A). (C) A mid-rooted phylogeny with three clades denoted 2.1 (grey, n=6), 2.2 (purple, n=5) and 2.3 (orange, n=3). Nodes with bootstrap support > 90 are shown. (D) PCA based on SNP data. (E) A network where each node represents one sample. Grey lines indicate inter-community connections, and black lines indicate intra-community ones.

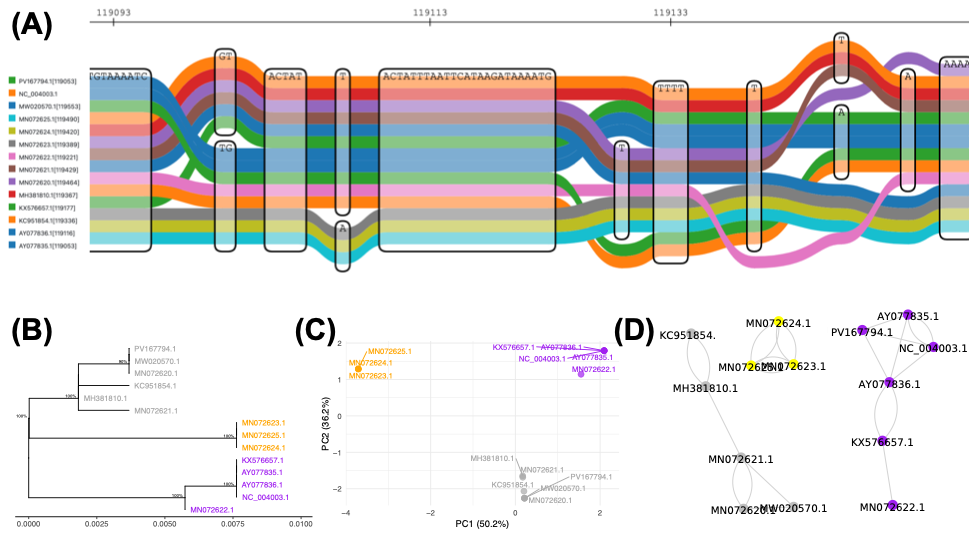

**Figure S16**. Visualisations of the region similar to LSDV132 (at NC_004003.1:1:119,053-119,613) gene in the GTPV PVG. (A) The sequenceTubeMap image shows an excerpt from the PVG at this region with with higher diversity in Clades 2.2 and 2.3. (B) A mid-rooted phylogeny based on SNPs with three clades denoted 2.1 (grey, n=6), 2.2 (purple, n=5) and 2.3 (orange, n=3). Nodes with bootstrap support > 90 are shown. (C) PCA based on SNP data. (D) A network from where each node represents one sample. Grey lines indicate inter-community connections, and black lines indicate intra-community ones.

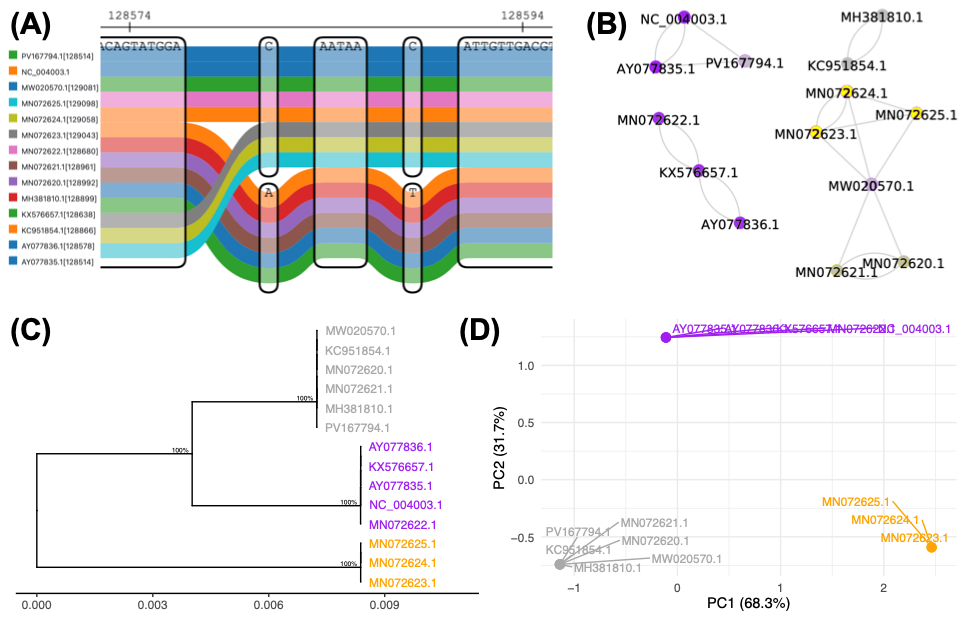

**Figure S17**. Visualisations of the region similar to LSDV136 (at NC_004003.1:128,526-128,998) gene in the GTPV PVG. (A) The sequenceTubeMap image shows an excerpt from the PVG at this region showing clade-specific genome graphs. (B) A network from where each node represents one sample. Grey lines indicate inter-community connections, and black lines indicate intra-community ones. (C) A mid-rooted phylogeny based on SNPs with three clades denoted 2.1 (grey, n=6), 2.2 (purple, n=5) and 2.3 (orange, n=3). Nodes with bootstrap support > 90 are shown. (D) PCA based on SNP data.

**Figure S18.** (A) The number of nonsynonymous SNPs per gene for SPPV (red) and GTPV (blue) (y-axis) versus LSDV (x-axis). (B) A midpoint-rooted phylogeny for gene LSDV039 showing LSDV (black), SPPV (red) and GTPV (blue).

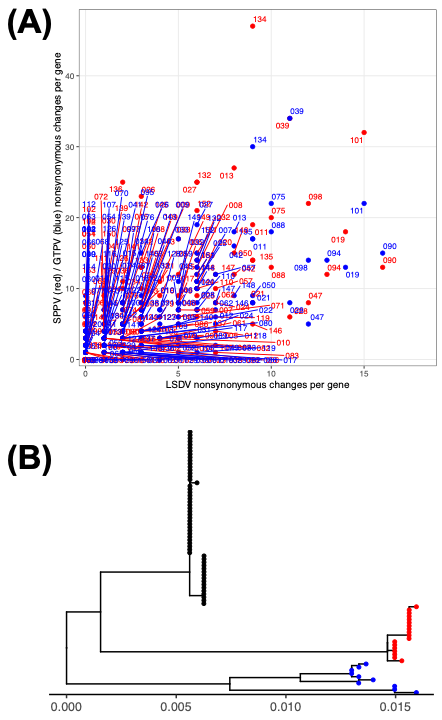
